# ACSL4 activity drives TNBC metastasis by positively regulating Histone H3 Acetylation mediated SNAIL expression

**DOI:** 10.1101/2023.10.16.562466

**Authors:** Abhipsa Sinha, Krishan Kumar Saini, Kiran Tripathi, Muqtada Ali Khan, Saumya Ranjan Satrusal, Ayushi Verma, Biswajit Mandal, Priyanka Rai, Sanjeev Meena, Mushtaq Ahmad Nengroo, Manish Pratap Singh, Namratha Shashi Bhushan, Madavan Vasudevan, Atin Singhai, Kulranjan Singh, Anand Kumar Mishra, Dipak Datta

**Author notes:** **Corresponding Author:** Dipak Datta, Division of Cancer Biology, CSIR-CDRI, B.S. 10/1, Sector 10, Jankipuram Extension, Sitapur Road, Lucknow-226031, India, (Extn-4347/48).

## Abstract

Triple-Negative Breast Cancer (TNBC) has profound unmet medical need globally for its devastating clinical outcome associated with rapid metastasis and lack of targeted therapies. Recently, lipid metabolic reprogramming has emerged as a major driver of breast cancer metastasis. Here, we unveil a strong association between the heightened expression of fatty acid metabolic enzyme, acyl-CoA synthetase 4 (ACSL4) and TNBC, which is primarily attributed by the selective absence of progesterone receptor (PR). Loss of ACSL4 function, either through genetic ablation or pharmacological inhibition significantly reduces metastatic potential of TNBC. Global transcriptome analysis reveals that ACSL4 activity markedly influences the gene expression pattern associated with TNBC migration. Mechanistically, ACSL4 alters fatty acid oxidation (FAO) and cellular acetyl-CoA levels, leading to the hyper-acetylation of particularly H3K27Ac and H3K9Ac marks resulting in overexpression of SNAIL during the course of TNBC metastatic spread to lymph node and lungs. Further, human TNBC metastasis exhibits positive correlation between ACSL4 and SNAIL expression. Altogether, our findings provide new molecular insights regarding the intricate interplay between metabolic alterations and epigenetic modifications, intertwined to orchestrate TNBC metastasis and posit a rational understanding for the development of ACSL4 inhibitors as a targeted therapy against TNBC.

## Introduction

According to a global cancer statistics report in 2020, breast cancer ranked the most commonly diagnosed cancer in women, accounting for 11.7% of all cancer instances, with an estimated 2.3 million new cases. Breast cancer is the fifth leading cause of cancer worldwide, with 685,000 deaths ^1^. Triple-negative breast cancer (TNBC), lacks estrogen receptor (ER), progesterone receptor (PR), and human epidermal growth factor receptor 2 (HER2) expressions ^2–5^ and is known to be the most aggressive subtype of breast cancer with a poor prognosis and adverse clinical outcomes ^6^. Early metastasis is responsible for more than 90% TNBC patient mortality and morbidity ^7, 8^. This underscores the urgent need to explore more effective therapeutic strategies for TNBC, particularly focusing on metastasis.

Metabolic reprogramming is one of the fundamental hallmarks of cancer ^9, 10^. More recently, a high resolution metastasis map has unveiled lipid metabolic reprogramming as a crucial hallmark of basal-like breast cancer or TNBC ^11^. Fatty acid oxidation (FAO) has lately been implicated in the orchestration of metastatic cancer progression ^12–14^ . Acyl-CoA synthetase 4 (ACSL4) is one of the five isoforms of mammalian long chain acyl-CoA synthetases that convert free long chain fatty acids (FAs) to fatty acyl-CoA esters, which then serves as a substrate for FAO ^15^. ACSL enzymes are characterized by their unique substrate and tissue specificity and play an integral role in the metabolic reprograming of cancer cells ^16, 17^. ACSL4 has been identified as an important biomarker for ferroptosis ^18–20^. While some reports suggest a conspicuous upregulation of ACSL4 in colon, hepatocellular, and aggressive breast carcinoma, where it has convincingly shown to play a pivotal role in driving increased proliferation, migration, and invasion in these malignancies ^21–24^, the biological implications of ACSL4 in advancing cancer metastasis remain enigmatic and have not yet been fully elucidated.

One of the major products of FAO is acetyl-CoA, which serves as a central metabolic intermediate and functions as a key signaling molecule promoting cell growth and proliferation ^25, 26^. In addition to being essential for maintaining cellular energy levels, acetyl-CoA is an important substrate for protein lysine acetylation, best known for regulating gene expression through histone acetylation^27^. Epigenetic changes are gaining immense prominence as pivotal events in breast cancer progression and metastasis^28^. Our recent findings have revealed the role of epigenetic modulator EZH2 in reshaping the metastatic landscape of TNBC ^29^. Additionally, there has been a growing comprehension regarding the crosstalk between metabolic reprogramming and epigenetic mechanisms during the course of cancer progression, epithelial to mesenchymal transition (EMT), and its subsequent metastasis ^30, 31^.

SNAIL (encoded by *SNAI1*) is a key transcription factor promoting EMT and metastatic dissemination ^32^. In TNBC, SNAIL is associated with early relapse, chemoresistance, and metastasis ^33, 34^. Emerging research suggests that chromatin modifiers like lysine specific demethylase (LSD1), Suv39H1 histone methyl transferase, and BRD4 contribute to the epigenetic regulation of SNAIL expression in varied cancer types, including TNBC ^35–37^. However, further investigation is required to gain a comprehensive understanding of epigenetic regulation of SNAIL and its functional implications, especially in the context of TNBC metastasis.

In this study, we unveil ACSL4 overexpression is strongly associated with TNBC subtype, where the absence of progesterone receptor (PR) is a critical factor contributing to elevated ACSL4 expression. Notably, the genetic silencing of ACSL4 reduces both tumor progression and metastasis; however, the pharmacological inhibition of its enzymatic activity predominantly impedes metastasis in cellular and mouse models of TNBC. Furthermore, our findings demonstrate that ACSL4 activity significantly influences TNBC metastasis by regulating SNAIL expression via modulation of histone H3 acetylation, mediated by altered FAO and cellular acetyl-CoA levels. Finally, we investigate the clinical relevance of our study in human TNBC metastasis, where a robust positive correlation was observed between ACSL4 and SNAIL expression compared to primary tumor. Taken together, these findings establish the role of ACSL4 as a key driver of TNBC metastasis and provide a rationale for the advancement of inhibitors that specifically target ACSL4 enzyme activity, thereby presenting a promising avenue for combatting TNBC metastasis.

## Results

### ACSL4 is selectively overexpressed in triple-negative breast cancer (TNBC)

Diverse gene expression studies have highlighted significant differences in the expression of lipid metabolic enzymes across different tumor types and molecular subtypes ^38–40^. The comparative assessment of mRNA expression patterns have elucidated noteworthy variations between receptor-positive (Non-TNBC) and triple-negative breast cancer (TNBC) in the context of lipid acquisition, storage and oxidation processes ^38, 41^. Although previous studies have shown the impact of lipogenesis with respect to dysregulation of lipid metabolism in cancer, the relevance of FAO for cancer cell function has not been well explored and its significance has remained obscure ^42^. To determine whether FAO is deregulated in TNBC, we analyzed the expression of the most relevant genes known to regulate the FAO pathway. Transcriptomic analyses of 514 breast cancer patients from the TCGA (BRCA) dataset revealed that out of the genes examined *ACSL4* exhibits a significant differential expression, notably distinguishing between Non-TNBC and TNBC subtypes (Figure 1A). Specifically, we observed a marked elevated expression of *ACSL4* in TNBC compared to Non-TNBC cell lines (Figure 1B). We also found significantly higher ACSL4 protein levels in TNBC compared to Non-TNBC cells, which was consistent with our mRNA expression findings (Figure 1C). To understand the clinical relevance of our observations, we queried the mRNA expression of *ACSL4* in human Non-TNBC and TNBC tumor samples. Notably, *ACSL4* expression was remarkably upregulated in TNBC tumors compared to Non-TNBC tumor tissues (Figure 1D). Further, to establish the above correlation in-situ within tumor tissues, we analysed ACSL4 protein expression in multiple human Non-TNBC and TNBC tumor samples. IHC staining of ACSL4 and subsequent H-scoring in these tumor samples clearly demonstrate a significant elevation of ACSL4 expression in TNBC tumor tissues as compared to Non-TNBC samples (Figure 1E and 1F). Together, our in-silico, in vitro and in-situ observations in human patients display a marked elevated expression of ACSL4 particularly in TNBC subtype as compared to Non-TNBC.

**Figure 1.**
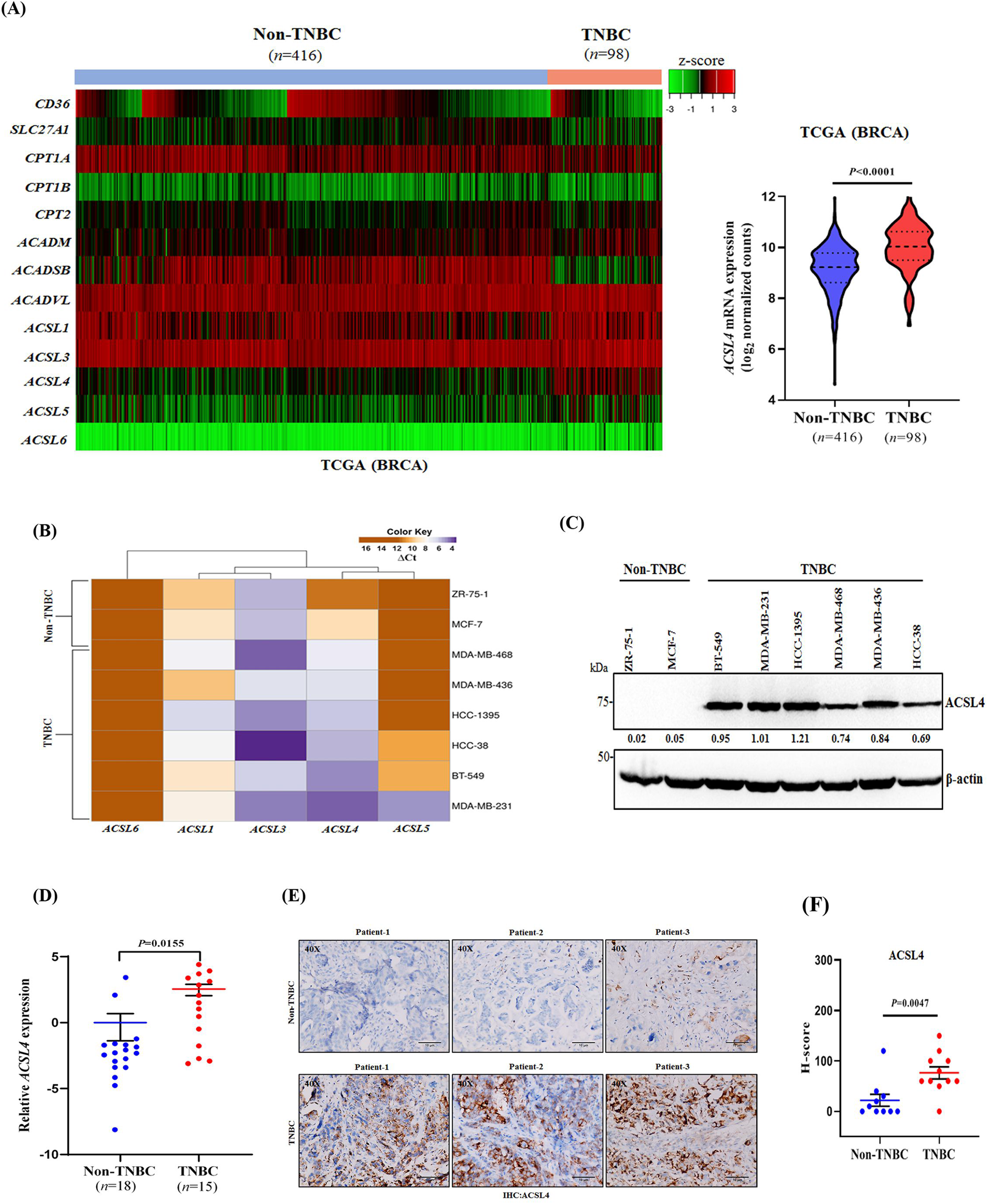
ACSL4 is selectively upregulated in TNBC subtype. **A.** Heat map depicting mRNA expression analysis of the key components of the FAO pathway in human Non-TNBC and TNBC tumors from the TCGA Breast Cancer (BRCA) dataset, which is publicly accessible via UCSC Xena Browser (left panel). The volcano plot represents the relative *ACSL4* expression between Non-TNBC and TNBC tumors according to the TCGA dataset (right panel). **B**. mRNA expression analysis of different *ACSL* isoforms in a panel of human breast cancer cell lines. The color key at the top indicates high (in purple) and low (in brown) ΔCt values in which ΔCt is the difference between Ct value of the target gene minus Ct value of the reference gene. **C**. Western blot analysis of ACSL4 protein levels in a panel of human breast cancer cell lines. β-actin was used as a loading control. The densitometric quantifications are provided at the bottom of each immunoblot. **D**. Analysis of *ACSL4* expression in TNBC compared to Non-TNBC patient tumor tissue samples. The dot plot represents the fold change values of *ACSL4* mRNA expression, data represent mean ± SEM, *n*=15 (TNBC) and *n=*18 (Non-TNBC) patient tumor tissue samples. **E.** Immunohistochemical staining of formalin-fixed paraffin embedded sections of human Non-TNBC and TNBC primary tumors using anti-ACSL4 antibody. Representative photomicrographs were shown at 40X magnification. Scale bar, 10 μm (40×). **F.** Quantitative H-scores for Non-TNBC (*n=*10) and TNBC (*n* =11) primary tumors were calculated for ACSL4 expression and represented as dot plot; error bar, mean ±SEM. For panels, **A** (left panel)**, D** Unpaired two-tailed Welch’s *t*-test and **F** Student *t*-test (two-tailed) were applied.

### Lack of Progesterone Receptor (PR) in TNBC promotes ACSL4 overexpression

Several observational and in-silico studies have demonstrated an inverse relationship between *ACSL4* mRNA expression and the expression of ER, PR, and HER2 in both breast cancer cell lines and tumor tissues ^43^. Furthermore, *ACSL4* mRNA expression is highest in triple-negative breast tumors (TNBCs) with claudin-low and basal-like subtypes ^44^. To enhance our empirical comprehension and for a more scientific understanding of the mechanism by which hormone or growth factor receptors regulate the expression of *ACSL4* in breast cancer, we generated stable triple positive MCF-7 cell lines harboring *ESR1*, *PGR*, and *ERBB2* knockdown, respectively (Figure 2A). Interestingly, we observed that selectively *PGR* knockdown but not ER or HER2 led to a pronounced increase in the expression of ACSL4 (Figure 2B). To substantiate our assertions, we ectopically restored the expression of ERα, PR-B/A (two independent PR isoforms), and HER-2 in triple-negative MDA-MB-231 cells, respectively and observed that neither *ESR1* nor *ERBB2* overexpression, but rather *PGR* overexpression decreased ACSL4 protein levels in TNBC (Figure 2C). Furthermore, we have also determined that *PGR* knockdown as well as treatment of PR antagonist (RU486) triggered a marked induction of *ACSL4* mRNA expression in MCF-7 cells (Figure 2D and 2E). The role of PR has been extensively defined, encompassing both its role as a transcriptional enhancer ^45^ and as a transcriptional suppressor ^46^. Therefore, to investigate whether PR directly regulates *ACSL4* transcription, we evaluated the potential existence of PR binding sites within the *ACSL4* promoter region and scanned the region for the presence of progesterone response elements (PREs) or NR3C binding motifs by employing publicly available transcription factor binding prediction software, JASPAR (http://www.jaspar.genereg.net). By utilizing the publicly available software Eukaryotic Promoter Database (https://epd.epfl.ch/)), we searched for the PREs in the *ACSL*4 promoter to corroborate the sequence of the PR DNA binding motif. Several putative PREs within the ∼-1 kb region upstream of transcription start site (TSS) of *ACSL4* were identified (Figure 2F, top and bottom panels). To confirm PR binding on the *ACSL4* promoter, we performed ChIP-quantitative PCR (ChIP-qPCR) for PR in MCF-7 cells, and a significant enrichment for PR-binding at two distinct sites (-0.3 kb and -0.6 kb upstream of TSS) was observed (Figure 2G). Next, we performed ChIP-qPCR for PR in order to determine the transcriptional activity of *ACSL4* upon stimulation with RU486, a PR antagonist. Interestingly, a marked reduction in the occupancy of PR on the *ACSL4* promoter indicated its transcriptional activation in RU486 stimulated MCF-7 cells (Figure 2H). To confirm the PR mediated transcriptional repression of *ACSL4*, we performed luciferase reporter assay in *PGR* knockdown MCF-7 cells, transfected with *ACSL4* promoter luciferase construct (-1 kb-consisting both proximal and distal promoter regions) plasmid. A significant increase in the luciferase activity was observed in *PGR* knockdown cells in comparison to control (Figure 2I). Similarly, a marked increase in the luciferase activity was observed upon RU486 treatment (Figure 2J). Together these findings suggest that the PR acts as a direct transcriptional repressor of *ACSL4*, and attenuating PR transcriptional function either by genetic knockdown or by using PR-antagonist relieve *ACSL4* transcriptional repression resulting in its upregulation.

**Figure 2.**
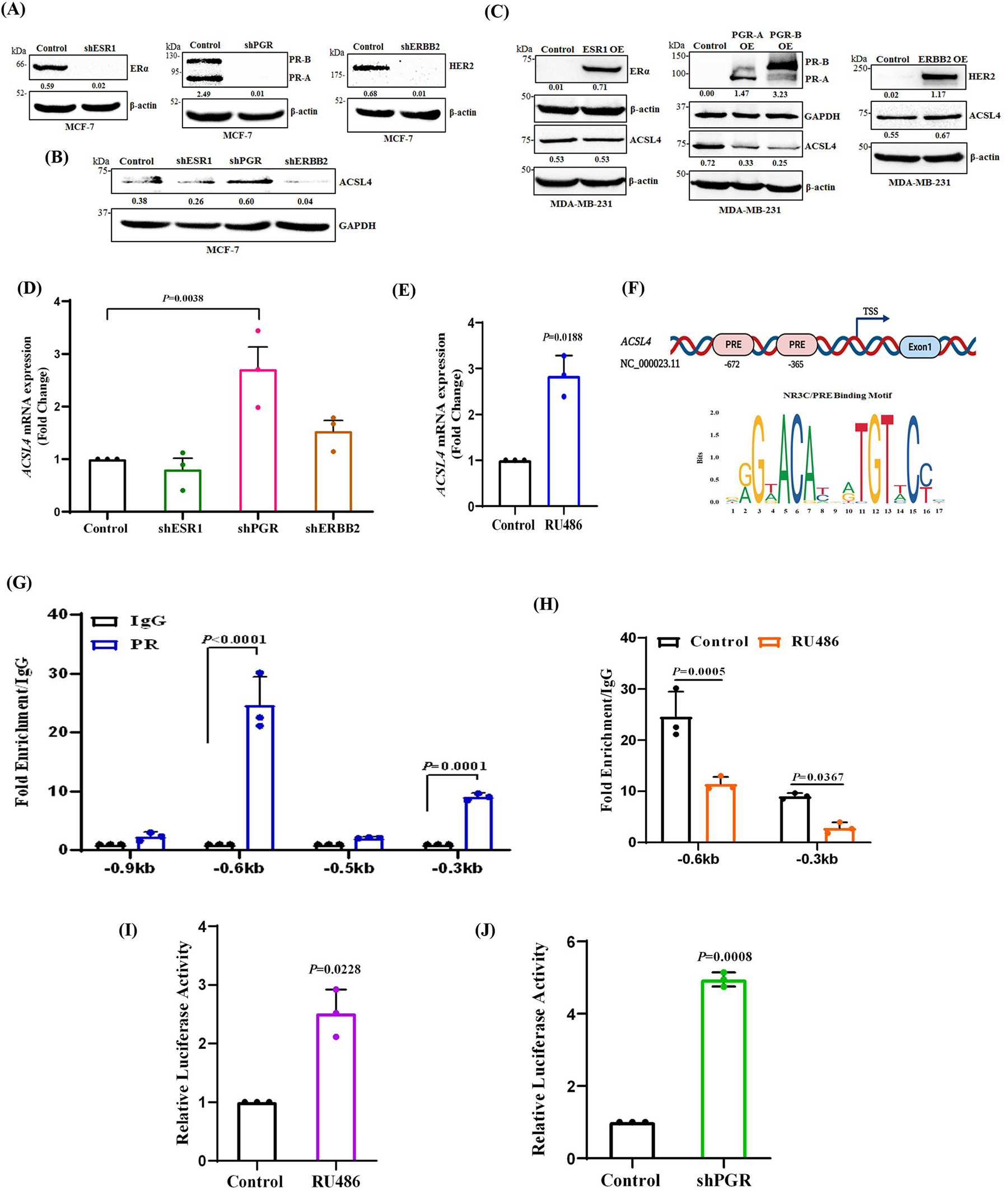
Progesterone Receptor (PR) directly binds to *ACSL4* promoter region and inversely regulates its expression. **A.** Immunoblot analysis of ERα, PR B/A, and HER-2 in control, *ESR1*, *PGR*, and *ERBB2* knockdown MCF-7 cells respectively. **B.** Immunoblot analysis of ACSL4 protein levels in control, *ESR1*, *PGR*, and *ERBB2* knockdown MCF-7 cells. **C**. Western blot analysis of ERα, PR B/A, HER-2, and ACSL4 in *ESR1* overexpression (OE), *PGR B/A* OE, and *ERBB2* OE MDA-MB-231 cells with their respective controls. **A-C**. β-actin and/or GAPDH were used as a loading control for the respective immunoblots. The densitometric quantifications (bottom of each immunoblot) are shown. **D.** mRNA expression analysis of *ACSL4* in *ESR1*, *PGR*, and *ERBB2* knockdown compared to control MCF-7 cells by RT-PCR. Data points are the mean of triplicate readings of samples; error bars, ±S.D. **E.** The MCF-7 cells were treated with either vehicle control or 20 μM of RU486 (PR antagonist) for 36 h followed by the analysis of *ACSL4* expression by qPCR. Data points are the mean of triplicate readings of samples; error bars, ±S.D. **F.** Diagrammatic scheme showing genomic location for the PREs on the *ACSL4* promoter (top). Bottom panel showing PR binding motif obtained from JASPAR database. **G**. ChIP-qPCR data in MCF-7 cells depicting recruitment of PR on the *ACSL4* promoter using primer pairs targeting −0.9, -0.6, -0.5, and, -0.3 kb of *ACSL4* gene. **H.** ChIP-qPCR data depicting the enrichment of PR on the *ACSL4* promoter at -0.3 kb and -0.6 kb regions upstream of TSS in MCF-7 cells either treated with vehicle control or RU486 (20 μM) for 36 h. **G, H** Each bar represents a mean of triplicate readings of fold enrichment normalized to IgG, error bars, ±S.D of. **I.** Luciferase reporter activity of the *ACSL4* promoter -1 kb region from TSS in control and PGR knockdown MCF-7 cells. **J.** Same as in **I** except either vehicle control or RU486 (20 μM) treated MCF-7 cells were used. **I, J** Each bar represents a mean of triplicate readings of samples, error bars, ±S.D. For panels, **D** One-way Anova Dunnett’s multiple comparisons test, **E, I, J** Unpaired two-tailed Welch’s *t*-test, **G, H** Two-way Anova Sidak’s multiple comparisons test were applied.

### *ACSL4* knock down significantly impedes TNBC growth and metastasis

ACSL4 was found to be selectively overexpressed in TNBC subtype and loss of PR could be one of the major reasons for the same. This result prompted us to further evaluate the role of ACSL4 in modulating TNBC tumor growth and metastasis. To address this, we generated stable *ACSL4* knockdown and control 4T-1 cells (Figure 3A). Though there was no notable differences in cell proliferation (Figure 3B), there was an evident decreased migratory potential of *ACSL4* knockdown cells compared to control cells as detected in wound healing assay (Figure 3C). Similarly, in trans-well chamber assay, *ACSL4* knockdown cells also show reduced invasive capabilities than the control cells (Figure 3D). The marked decrease in the migratory and invasive ability of *ACSL4* knockdown cells encouraged us to explore its further impact on TNBC metastasis. To investigate the metastatic potential of TNBC cells upon ablation of *ACSL4*, we employed bioluminescence imaging of nude mice following orthotopic inoculation of Luc-tagged control and *ACSL4* knockdown 4T-1 cells. Compared to control, depletion of ACSL4 results in significant reduction in tumor growth and metastasis (Figure 3E-3F). Additionally, bioluminescence imaging of harvested organs from control and *ACSL4* knockdown tumor bearing mice suggests that ACSL4 depletion greatly inhibits TNBC metastasis in the secondary organs like lung, liver and spleen (Figure 3G). To validate the significance of the observed phenotype regarding the association of reduced ACSL4 levels with delayed TNBC metastasis, we designed a robust experimental plan which is described in Figure 3H. Here, we orthotopically inoculated 4T-1 Td-tomato-tagged control cells and 4T-1 untagged *ACSL4* knockdown cells into the left and right mammary fat pad of nude mice, respectively. This was followed by isolation of tumor cells; 25 days post inoculation from distant metastatic sites. The FACS analysis of single cells isolated from the distinct metastatic sites (CTCs, lymph node, lung, liver and peritoneal fluid) of the experimental mice shows a robust anti-metastatic effect upon silencing *ACSL4* as we found a significantly less percentage of *ACSL4* knockdown cells as compared to control in a pool of 100% metastatic cells (Figure 3I and Supplementary Figure 1A). To validate our observation in human TNBC model, we initiated the silencing of individual *ACSL* family members (*ACSL1*, *ACSL3*, *ACSL4*, *ACSL5*, and *ACSL6*) in MDA-MB-231 cells and assess their migration potential. Strikingly, the knockdown of all *ACSL* isoforms (*ACSL1*, *ACSL3*, *ACSL5*, and *ACSL6*) except *ACSL4* yielded negligible effect on the migration of TNBC cells (Figure 3L and Supplementary Figure 2A-2E). In addition to findings presented above, we generated *ACSL4* knockdown in MDA-MB-231 cells (Figure 3J) and observed that the genetic ablation of *ACSL4* did not impair cell proliferation but substantially decreased TNBC migration and invasion (Figure 3K-3M). Next, we performed ACSL4 gain of function experiment by generating ACSL4 overexpressing stable TNBC cells (Figure 3N) and assess its proliferative (colony formation assay) and migratory potential (wound healing and invasion assay). As observed in Figure 3N-3P, *ACSL4* OE cells have markedly greater migratory potential, compared to control cells in vitro, whereas cell proliferation remains unaffected due to ACSL4 overexpression. Equivalent to the observations in 4T-1-Luc model, the MDA-MB-231-Chili-Luc-tagged *ACSL4* knockdown cells after orthotopic inoculation in NSG mice exhibited significant reduction in tumor growth as well as attenuated metastatic dissemination compared to the control group as evaluated by sequential bioluminescence imaging of tumor-bearing mice and harvested organs from respective groups (Figure 3Q-3S). Collectively, our observations strongly suggest that ACSL4 indeed plays a pivotal role in modulating TNBC growth and metastasis in vivo.

**Figure 3.**
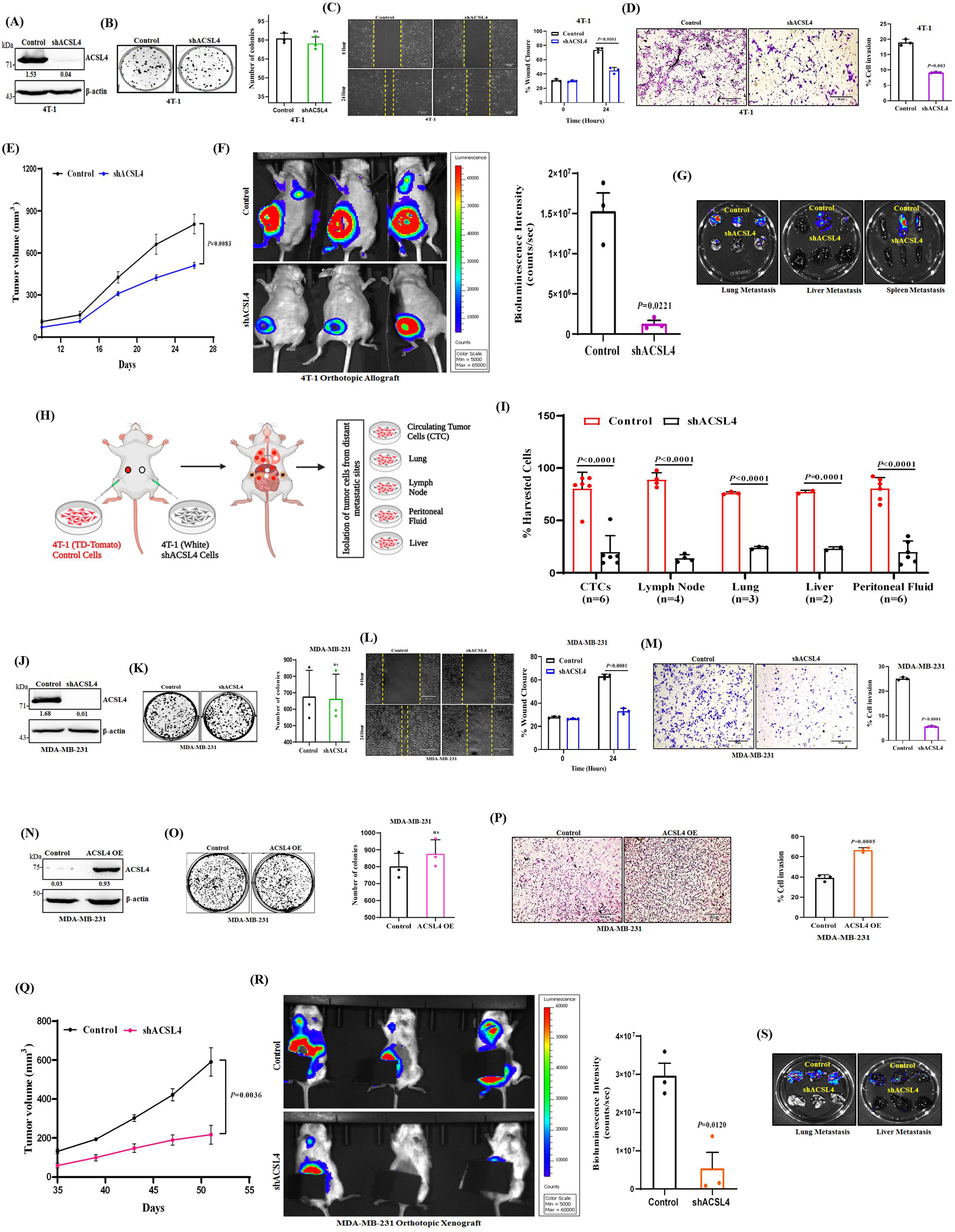
Genetic knockdown of *ACSL4* impairs TNBC growth and metastasis. **A.** The *ACSL4* knockdown was confirmed through immumoblot using shRNA mediated lentiviral transduction in 4T1 cells. β-actin was used as a loading control. **B**. Representative images of the colony formation assay in control and *ACSL4* knockdown 4T-1 cells (left panel). The quantitative analysis of the colony formation assay is provided (right panel). Each bar represents a mean of triplicate readings of samples, error bars, ±S.D. **C.** Representative images of the wound healing assay to measure the migration ability of control and *ACSL4* knockdown 4T-1 cells (left panel) at the indicated time points (0 and 24 h), magnification 10×. Scale bar 50 µm. The quantitative bar graphs of the wound healing assay is shown (right panel), representing a mean of triplicate readings of samples, error bars, ±S.D. **D.** Representative images of the trans-well chamber invasion assay to measure the invasive ability of control and *ACSL4* knockdown 4T-1 cells at 24 h (left panel), magnification 10×. Scale bar 50 µm. The quantitative bar graphs of trans-well chamber invasion assay is shown (right panel). Each bar represents a mean of triplicate readings of samples, error bars, ±S.D. **E.** Tumor growth curve of *ACSL4* knockdown and control 4T-1 cells orthotopically implanted in nude mice (*n*=7 per group), tumor volume was measured at regular intervals, data points are indicated as mean, error bar ± SEM. **F.** In-vivo bioluminescence images of tumor-bearing control (top panel), and *ACSL4* knockdown (bottom panel) mice. The color scale indicates the bioluminescence intensity (counts/sec) emitted from each group. Quantitative bar graph representation (right panel) of the metastatic burden in terms of bioluminescence intensity, calculated from the region of interest (ROI) is shown, (*n*=3) each group. Each bar represents triplicate readings of samples, mean ± SEM. **G.** Representative bioluminescence images of lung, liver, and spleen harvested from control and *ACSL4* knockdown tumor-bearing mice. **H.** The schematic representation of the experimental planning. **I.** The quantitative bar graph represents the flow cytometry analysis of the percentage of control (Td-Tomato^+^) and *ACSL4* knockdown (white) cells isolated from several metastatic sites (CTCs, lymph node, lung, liver, and peritoneal fluid), data are represented as mean, error bar ± SD. **J.** Western blot analysis of *ACSL4* knockdown in MDA-MB-231 cells. β-actin was used as a loading control. **K-M**. Representative images of the colony formation (**K**, left panel), wound healing (**L**, left panel), and trans-well chamber invasion assays (**M**, left panel) in control and *ACSL4* knockdown MDA-MB-231 cells, 10× magnification. Scale bar 50 µm. The quantitative analysis of the colony formation (**K,** right panel), wound healing assay (**L**, right panel), and trans-well (**M,** right panel) assay are shown. Each bar represents a mean of triplicate readings of samples, error bar, ±SD. **N.** Immunoblot analysis for the expression of ACSL4 in control and *ACSL4* overexpression (OE) in MDA-MB-231 cells. β-actin was used as a loading control **O.** Same as **K**, except *ACSL4* OE MDA-MB-231 cells was used. **P.** Same as **M,** except *ACSL4* OE MDA-MB-231 cells was used. **Q.** Tumor growth curve of *ACSL4* knockdown and control MDA-MB-231 cells orthotopically inoculated in NSG mice, (*n*=5) each group, tumor volume was measured at regular intervals, data points are indicated as mean, error bar ± SEM. **R.** Representative bioluminescence images of tumor-bearing control (top panel), and *ACSL4* knockdown (bottom panel) mice. The color scale indicates the bioluminescence intensity (counts/sec) emitted from each group. Quantitative bar graph representation (right panel) of the metastatic burden in terms of bioluminescence intensity, calculated from the region of interest (ROI) is shown (*n*=3) each group. Each bar represents triplicate readings of samples, mean ± SEM. **S.** The bioluminescence analysis of the metastatic signal in the organs harvested from control and *ACSL4* knockdown mice. For panels, **B**, **D, F, K, M, O**, **P**, **R** Unpaired two-tailed Welch’s *t*-test, **C**, **I**, **L** Two-way Anova Sidak’s multiple comparisons test, **E**, **Q** Student’s *t*-test were applied. The densitometric quantifications are provided at the bottom of each immunoblot.

### Inhibition of ACSL4 enzymatic activity perturbs TNBC metastasis

To dissect the contribution of ACSL4 protein versus its enzymatic activity in modulating TNBC metastasis, we made use of ACSL4 pharmacological inhibitor PRGL493 which strongly inhibits its enzymatic activity, without attenuating its expression ^47^. In this particular context, we conducted a thorough assessment of the potential of PRGL493 to impede TNBC migration, invasion, and metastasis in two (4T1 and MDA-MB-231) distinct TNBC models. As previously reported, we did not observe any discernible alteration in ACSL4 protein expression subsequent to PRGL493 treatment in 4T-1 cells (Figure 4A). While the colony formation assay revealed no significant changes in cellular proliferation (Figure. 4B), there was a marked impairment in the migratory and invasive capabilities of 4T1 cells in comparison to the control cells, as determined by using wound healing and trans-well chamber invasion assays, respectively (Figure 4C and 4D). Consistent with our in vitro findings, we observed no significant changes in the primary tumor burden but a notable reduction in metastasis following intra peritoneal administration of PRGL493 (500 µg/kg) in 4T1-Luc allograft TNBC orthotopic mice model between the control and treatment group. Furthermore, serial bioluminescence imaging and analysis of the harvested organs from control and PRGL493 treated tumor-bearing mice indicate that administration of PRGL493 considerably dampens TNBC metastasis (Figure 4E-4G). In extension to our preceding observations, we have replicated similar results in the context of MDA-MB-231 TNBC cells. As depicted in Figure 4H-4K, it is evident that PRGL493 has no visible impact on the protein expression levels of ACSL4. Moreover, its influence on the proliferation of MDA-MB-231 cells is negligible. However, a substantial disruption in relation to the migration and invasion potential of TNBC cells are observed due to the pronounced impact exerted by PRGL493. Next, we wish to explore the wider implications of our findings, in this endeavor we employed another preclinical metastatic TNBC orthoptopic xenograft model, where we inoculated human MDA-MB-231-Chili-Luc cells in NSG mice. Concordance to the results obtained in the 4T1-Luc model, we observe marked decrease in metastasis following intra-peritoneal dosing of PRGL493 (500 µg/kg), at consecutive days though the differences in the primary tumor volume remains insignificant between the control and treated group (Figure 4L-4N). Overall, our in vitro and in vivo data suggest that loss of enzymatic function of ACSL4 but not its protein expression resulted in significant inhibition of TNBC metastasis though primary tumor growth remains minimally affected.

**Figure 4.**
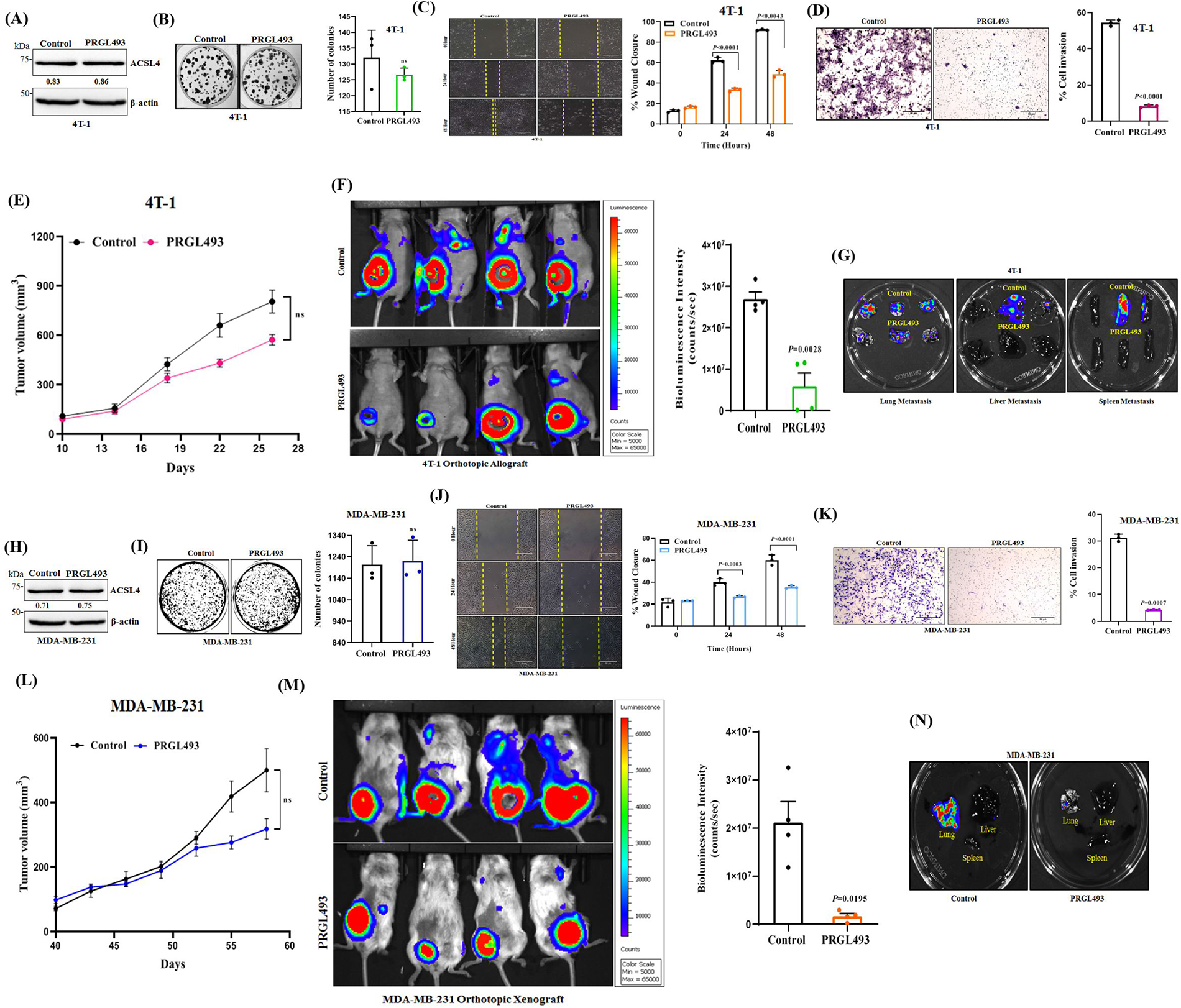
Inhibition of ACSL4 activity significantly impedes TNBC metastasis. **A.** Immunoblot analysis of ACSL4 proteins levels in 4T-1 cells either treated with vehicle control or 20 µM of PRGL493 (ACSL4 inhibitor). β-actin was used as a loading control. **B.** Representative images of the colony formation assay to access the proliferation ability of 4T-1 cells, treated with either vehicle control or PRGL493 (20 µM) (left panel). The quantitative analysis of the colony formation assay is provided (right panel). Each bar represents a mean of triplicate readings of samples, error bars, ±S.D. **C.** Representative images of the wound healing assay to measure the migration ability of 4T-1 cells, either treated with vehicle control or PRGL493 (20 µM) (left panel) at the indicated time points (0, 24 h, and 48 h), magnification 10×. Scale bar 50 µm. The quantitative bar graphs of the wound healing assay is shown (right panel), representing a mean of triplicate readings of samples, error bars, ±S.D. **D.** Representative images of the trans-well chamber invasion assay to measure the invasive ability of 4T-1 cells, either treated with vehicle control or PRGL493 (20 µM) at 24 h (left panel), magnification 10×. Scale bar 50 µm. The quantitative bar graphs of trans-well chamber invasion assay is shown (right panel). Each bar represent a mean of triplicate readings of samples, error bars, ±S.D. **E.** Tumor growth curve of 4T-1 TNBC orthotopic tumor allograft in nude mice, treated either with vehicle or PRGL493 (500 μg/kg/every other day). Tumor volume was measured at regular intervals and mice were randomised and separated for treatment when tumor volume reaches 100 mm^3^ in size. Data represent the means of tumor volume ± SEM, (*n=*7) each group. **F.** Representative bioluminescence images of 4T-1 tumor-bearing mice, treated with either vehicle (top panel) or PRGL493, bottom panel. The color scale indicates the bioluminescence intensity (counts/sec) emitted from each group. Quantitative bar graph representation (right panel) of the metastatic burden in terms of bioluminescence intensity, calculated from the region of interest (ROI) is shown, (*n*=4) each group. Each bar represents quadruplicate readings of samples, mean ± SEM. **G.** Representative bioluminescence images of lung, liver, and spleen harvested from control and PRGL493 treated tumor-bearing mice. **H.** Immunoblot analysis of ACSL4 proteins levels in MDA-MB-231 cells either treated with vehicle control or PRGL493 (20 µM) for 48 hours. β-actin was used as a loading control. **I-K**. Representative images of the colony formation (**I**, left panel), wound healing (**J**, left panel), and trans-well chamber invasion assays (**K**, left panel) in control and PRGL493 (20 µM) treated MDA-MB-231 cells, 10× magnification. Scale bar 50 µm. The quantitative analysis of the colony formation (**I,** right panel), wound healing assay (**J**, right panel), and trans-well (**K,** right panel) assay are shown. Each bar represents a mean of triplicate readings of samples, error bar, ±SD. **L.** Tumor growth curve of NSG mice bearing MDA-MB-231 breast tumor xenograft, (*n*=6) each group, either treated with vehicle or PRGL493 (500 μg/Kg). Tumor volume was measured at the point when it reached approximately 100 mm^3^ in size. Data are represented as mean, error bar ± SEM. **M.** Representative bioluminescence images of MDA-MB-231 tumor-bearing mice, treated with either vehicle (top panel) or PRGL493 (500 μg/kg/every other day), bottom panel. The color scale indicates the bioluminescence intensity (counts/sec) emitted from each group. Quantitative bar graph representation (right panel) of the metastatic burden in terms of bioluminescence intensity, calculated from the region of interest (ROI) is shown, (*n*=4) each group. Each bar represents quadruplicate readings of samples, mean ± SEM. **N.** The bioluminescence analysis of the metastatic signal in the organs harvested from control and PRGL493 treated tumor-bearing mice. For panels, **B, D, F, I, K**, **M** Unpaired two-tailed Welch’s *t*-test, **C, J** Two-way Anova Sidak’s multiple comparisons test, **E, L** Student’s *t*-test were applied. The densitometric quantifications are provided at the bottom of each immunoblot.

### Transcriptome analysis reveals that ACSL4 activity alters gene expression pattern relevant to TNBC migration and invasion

Next, we aimed to unravel the molecular mechanisms mediated by ACSL4 in modulating TNBC metastasis. Here, we conducted a genome-wide analysis of transcript levels (RNA-sequencing [RNA-seq]) on MDA-MB-231 cells with three biological replicates in each group: control, sh*ACSL4* (*ACSL4* knockdown), and PRGL493 (inhibitor treatment), respectively. Following total RNA collection, we performed next generation sequencing to identify differentially expressed genes (DEGs) between control and *ACSL4* knockdown groups, as well as control and PRGL493 treated group (Figure 5A). The volcano plots (Figure 5B, top and bottom panels) and the heat map analysis results (Figure 5C, left and right panels) revealed that a total of 285 upregulated and 455 downregulated genes were identified in *ACSL4* knockdown cells compared to control and in case of PRGL493 treatment vs. control, a total of 363 upregulated and 884 downregulated genes were identified. To gain insights into the biological pathways perturbed upon loss of ACSL4 function, we focused on the common genes that were differentially expressed in the following two cases: (1) sh*ACSL4* vs. control and (2) PRGL493 vs. control (Figure 5D). Gene ontology enrichment analysis of 312 common DEGs identified in both cases revealed that the genes found to be robustly downregulated upon *ACSL4* ablation and due to its functional inhibition are predominantly involved in the regulation cell migration, cell motility, epithelial to mesenchymal transition, as well as other pathways that are critical for regulating cancer metastasis (Figure 5E). The heat map depicts the top 23 genes (Log_2_FC < -1 and *P* < 0.05) that have been identified to be downregulated in our RNA-seq analysis and are known to have critical roles in governing cell migration and motility (Figure 5F). We then validated top 25% of the downregulated genes, which included *HES1*, *EGR3*, *NR4A1, DUSP1, ARC, SNAI1, CXCL2,* and *NR4A2* in relation to the phenotypes observed in our study via RT-PCR upon *ACSL4* knockdown and inhibition of its activity by PRGL493, respectively (Figure 5G and 5H). Collectively, these analyses depict that ACSL4 activity considerably influences the gene expression pattern in TNBCs and plays an essential role in modulating the gene expression landscape of TNBC cell migration and metastasis.

**Figure 5.**
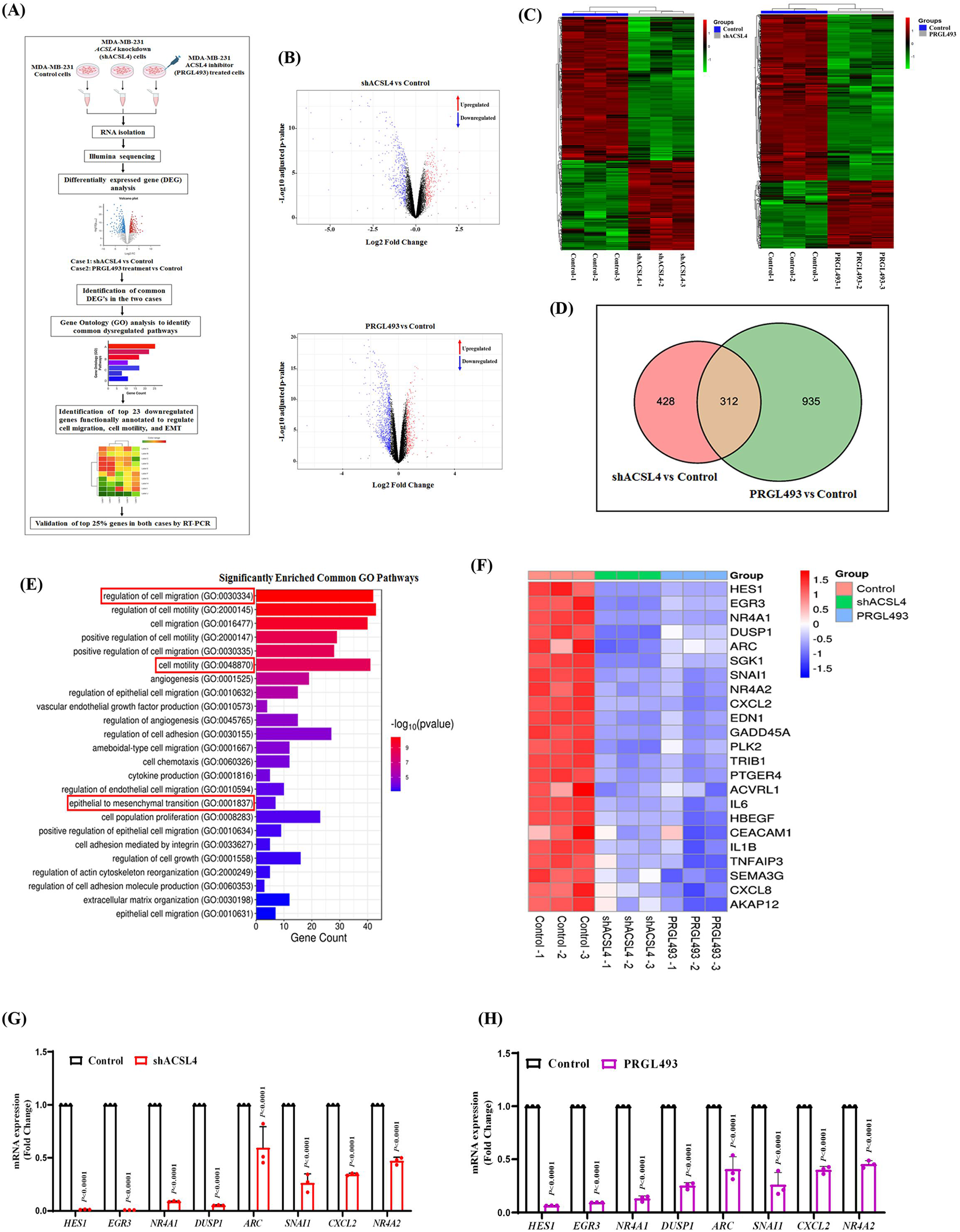
Transcriptome analysis revealed that loss of ACSL4 activity markedly inhibits pathways central to TNBC migration and invasion. **A.** Schematic representing the key steps involved in assessing the impact of loss of ACSL4 activity on the global transcription network through high-throughput transcriptome analysis with RNA isolated from control, *ACSL4* knockdown, and PRGL493 (20 µM)-treated MDA-MB-231 cells (*n*=3 in each case). **B.** Differential gene expression pattern upon gene knockdown and treatment compared to control, depicted in volcano plots-sh*ACSL4* vs control top panel) and PRGL493 vs control (bottom panel). The downregulated and upregulated genes are shown in blue and red, (Fold change (FC) >1.5 for upregulated and FC< 1.5 for downregulated genes, *P < 0.05*), respectively. **C.** Heat maps showing the differential gene expression pattern in *ACSL4* knockdown and PRGL493 treated cells compared to control [Color scale: green (downregulated), black (no change), and red (upregulated)]. **D.** Venn diagram depicting the number of differentially expressed genes upon *ACSL4* knockdown and PRGL493 treatment with respect to control. **E.** Gene Ontology Enrichment Analysis was performed using the 312 common DEGs that were altered in both *ACSL4* knockdown and PRGL493 treated conditions compared to control to investigate the different biological processes that were affected, as predicted by Panther software. **F.** Heat map shows differential expression of top 23 downregulated genes that are related to the regulation of cell migration, cell motility, and EMT. Expression is represented as a plot of normalized read counts for control, *ACSL4* knockdown, and PRGL493 treatment (*n* = 3 per group). **G.** Real-time qPCR validation of top 25% of downregulated cell migration-related genes identified by RNA-seq analysis in *ACSL4* knockdown compared to control. Each bar represents a mean of triplicate readings of samples; error bars, ±S.D. **H.** Same as **G**, except PRGL493 treatment was compared to control. For panels, **G** and **H** Two-way Anova Sidak’s multiple comparisons test was applied.

### ACSL4 activity governs histone H3 acetylation in TNBC by modulating Fatty Acid Oxidation

Notably, several studies have yielded intriguing findings with respect to deregulation of lipid metabolism in TNBC either by downregulation of activators of lipid biosynthesis along with enhanced expression of activators of Fatty Acid Oxidation (FAO) ^48–50^. In this context, we next sought to determine the impact of loss in ACSL4 activity on FAO, either by genetic ablation of *ACSL4* or treatment of MDA-MB-231 cells with PRGL493. First, we analysed ACSL activity in *ACSL4* knockdown and PRGL493 treated MDA-MB-231 cells. In both cases, we observed a marked decrease in ACSL activity compared to control and vehicle-treated cells (Figure 6A and 6B). Second, to assess the contribution of ACSL4 in modulating FAO, we measured OCR (Oxygen Consumption Rate) upon *ACSL4* silencing. Our observations indicated that the OCR rate derived from long chain fatty acids was significantly reduced in *ACSL4* knockdown cells compared to control cells (Figure 6C). Apart from genetic silencing of *ACSL4*, we employed treatment of MDA-MB-231 cells with PRGL493 to assess the impact of loss in ACSL4 activity on FAO. Here, etomoxir (a well-established FAO inhibitor) treated MDA-MB-231 cells was used as a positive control to validate the efficiency of PRGL493 to inhibit FAO. Interestingly, we observed that loss in ACSL4 activity by PRGL493 significantly reduces the OCR compared to vehicle-treated cells, indicating that FAO rate decreases substantially upon hindering ACSL4 activity (Figure 6D). FAO generates acetyl-CoA, a major substrate for histone acetylation. The considerable reduction in FAO upon inhibition of ACSL4 activity intrigued us to estimate cellular acetyl-CoA levels upon silencing of *ACSL4* or after treatment with PRGL493. As shown in (Figure 6E and 6F), the acetyl-CoA levels were significantly decreased in both *ACSL4* silenced and PRGL493 treated cells. Based on our preceding result that demonstrated the regulatory influence of ACSL4 on gene expression in relation to the decreased acetyl-CoA levels, we speculated that this association might be mediated by epigenetic mechanisms that govern histone acetylation processes in TNBC. To understand the effect of ACSL4 activity on histone acetylation we undertook different approaches. We first analysed the perturbation of acetylated histone marks upon genetic silencing of *ACSL4*. Acetylation of H3K9, H3K27, H3K14, H3K23, and H3K18 histone marks, which control active gene transcription, was decreased upon *ACSL4* knockdown (Figure 6G). Next, we evaluated the expression of histone acetylation marks upon overexpression of *ACSL4* and observed a marked upregulation of H3K9, H3K27, H3K14, and H3K23ac levels (Figure 6H). Further to investigate the role of ACSL4 activity in modulating histone acetylation, we treated MDA-MB-231 cells with PRGL493 and detected significant reduction in acetylated H3K9, H3K27, H3K14, H3K23, and H3K18 histone marks (Figure 6I). To demonstrate that the alterations in the expression of acetylated histone marks was the result of FAO inhibition by decreased ACSL4 activity, which led to the reduction of acetyl-CoA, we supplemented control and *ACSL4* knockdown cells with synthetic acetyl-CoA. This rescued the suppression of acetylation levels of H3K9, H3K27, H3K14, H3K23 histone marks in *ACSL4* knockdown cells (Figure 6J). Additionally, to characterize the changes in TNBC migration, we treated *ACSL4* knockdown cells with either with vehicle or synthetic acetyl-CoA. We found that exogenous acetyl-CoA supplementation rapidly increased the migration and invasion capabilities of *ACSL4* knockdown cells as demonstrated by wound healing and trans-well chamber invasion assays, respectively (Figure 6K and 6L). Taken together, our data convincingly suggests that ACSL4 plays a pivotal role in modulating FAO and predominately governs the acetylation of H3K9, H3K27, H3K14, and H3K23 histone marks, thereby connecting ACSL4 activity to the epigenetic regulation of genes involved in TNBC migration.

**Figure 6.**
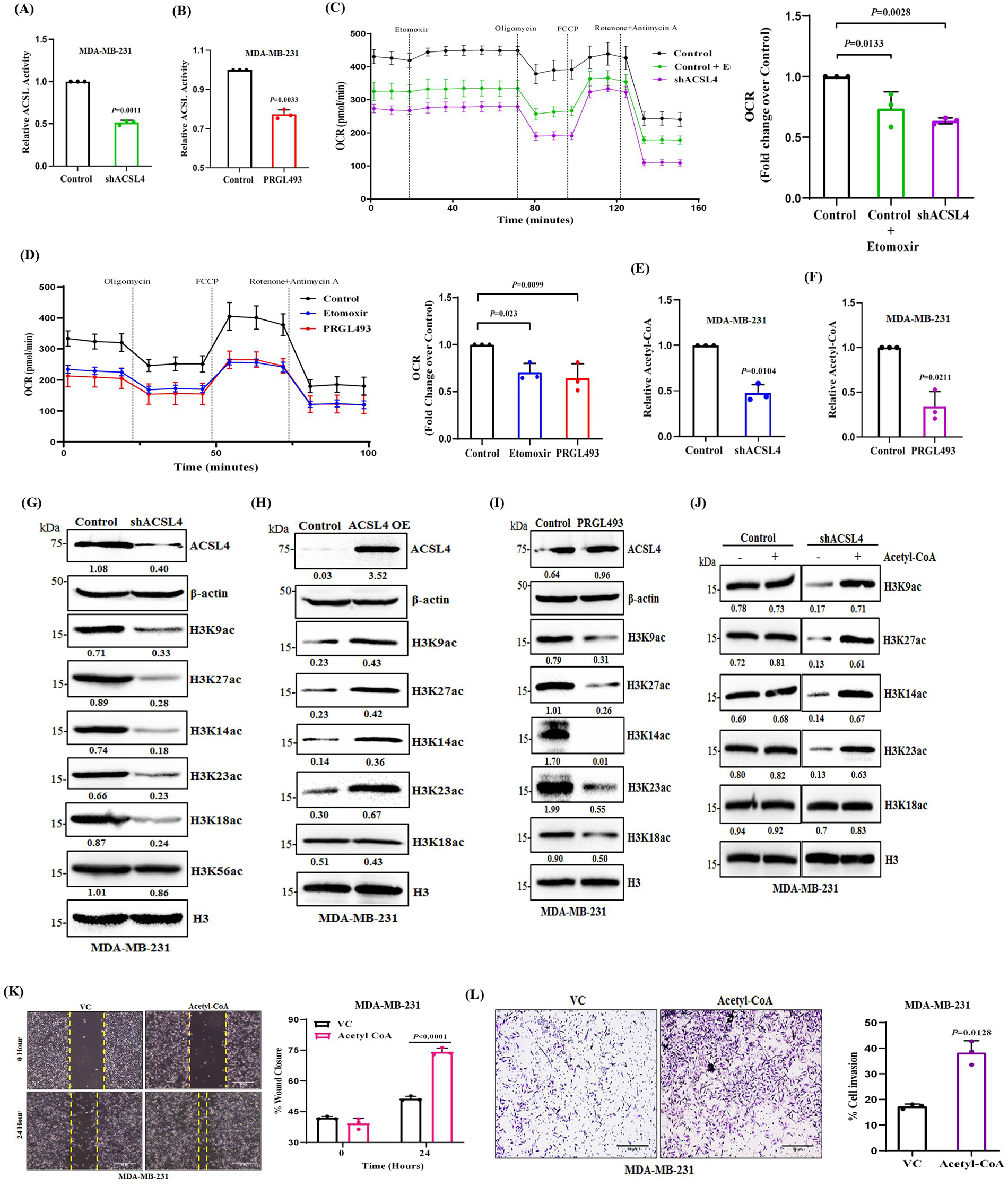
Impairment of ACSL4 activity reduces Fatty Acid Oxidation (FAO) and modulates histone H3 acetylation. **A** and **B.** Estimation of ACSL activity using acyl-CoA synthetase assay kit following manufacturer’s instruction. **A.** ACSL activity was assessed in *ACSL4* knockdown MDA-MB-231 cells compared to control **B.** ACSL activity was determined in MDA-MB-231 cells treated either with vehicle control or PRGL493 (20 µM) for 48 h. Each bar represents a mean of triplicate readings of samples; error bars, ±S.D. **C.** FAO was assessed by OCR (Oxygen Consumption Rate) in MDA-MB-231 cells using Agilent Seahorse XF Long Chain Fatty Acid Oxidation Stress Test. Specific inhibitors were added as indicated: Etomoxir (FAO inhibitor, blocks the transport of FAs into the mitochondria), Oligomycin (ATP synthase inhibitor), FCCP (carbonyl cyanide-4-phenylhydrazone, ATP synthesis uncoupler), Rotenone + Antimycin A (shuts down mitochondrial respiration by inhibiting complex I and complex Ш, respectively). Left panel, representative OCR profile in *ACSL4* knockdown cells compared to control. Right panel, quantification of OCR in *ACSL4* knockdown cells with respect to control. Each bar represents a mean of triplicate readings of samples; error bars, ±S.D. **D.** Left panel, representative oxygen consumption rate (OCR) profile in Etomoxir (10 µM) or PRGL493 (20 µM) treated MDA-MB231 cells (24h) compared to control, respectively. Right panel, quantitative bar graph representation of OCR. Each bar represents a mean of triplicate readings of samples; error bars, ±S.D. **E** and **F.** The acetyl-CoA production was detected using acetyl-CoA assay kit following the manufacturer’s instruction. **E.** The quantitative bar graph representation of acetyl-CoA levels in *ACSL4* knockdown MDA-MB231 cells compared to control. **F.** Same as **E**, except MDA-MB-231 cells treated either with vehicle control or PRGL493 (20 µM) for 48 h. Each bar represents a mean of triplicate readings of samples; error bars, ±S.D. **G-J.** Immunoblot analysis of different histone H3 acetylation marks in MDA-MB-231 cells by western blotting. β-actin and total H3 were used as loading control. The densitometric quantifications are shown at the bottom of each immunoblot. **G.** H3K9, H3K27, H3K14, H3K23, H3K18, and H3K56 acetylation levels in Control and *ACSL4* knockdown cells. H3K9, H3K27, H3K14, H3K23, and H3K18 acetylation levels in Control and *ACSL4* OE cells **(H),** and Control and PRGL493 (20 µM) for 48 h treated cells **(I). J.** The histone acetylation levels were assessed in the presence or absence of synthetic acetyl-CoA (acetyl-CoA trisodium salt) (20 µM) in control and *ACSL4* knockdown MDA-MB-231 cells **K.** Representative images of the wound healing assay to measure the migration ability of *ACSL4* knockdown MDA-MB231 cells after supplementation of synthetic acetyl-CoA (20 µM) (left panel), magnification 10×. Scale bar 50 µm. The quantitative bar graphs of the wound healing assay is shown (right panel), representing a mean of triplicate readings of samples, error bars, ±S.D. **L.** Representative images of the trans-well chamber invasion assay to measure the invasive ability of *ACSL4* knockdown MDA-MB231 cells after supplementation of synthetic acetyl-CoA (20 µM) (left panel), magnification 10×. Scale bar 50 µm. The quantitative bar graph (right panel) represents a mean of triplicate readings of samples, error bars, ±S.D. For panels, **A**, **B**, **E**, **F, L** Unpaired two-tailed Welch’s *t*-test, **C**, **D** (right panel) One-way Anova Dunnett’s multiple comparisons test, **K** Two-way Anova Sidak’s multiple comparisons test were applied.

### ACSL4 orchestrates histone H3 acetylation and positively regulates the modulation of *SNAI1* expression in TNBC

As described above, we demonstrated that ACSL4 activity modulates FAO and cellular acetyl-CoA levels. These alterations subsequently affect histone acetylation and supplementing acetyl-CoA reverses the impaired migration in TNBC caused by *ACSL4* ablation. To gain further mechanistic insights into the contribution of these changes in ascertaining TNBC migration and metastasis, we examined the mRNA expression of genes, as previously analysed in our RNA-seq analysis, after exogenous supplementation of acetyl-CoA in *ACSL4* silenced MDA-MB-231 cells. As shown in (Figure 7A), besides *SNAI1*, no other genes exhibited robust upregulation upon acetyl-CoA supplementation in *ACSL4* knockdown cells compared to control. Since SNAIL has been extensively and conclusively characterized as a master regulator of EMT process, initiating cancer cell plasticity, migration, and metastatic dissemination across diverse tumor types ^32, 51, 52^. The downregulation of SNAIL in *ACSL4* knockdown cells highlights its role in impeding the metastatic potential of TNBC, emphasizing the significance of lower SNAIL expression following ACSL4 activity loss. Next, we further investigated whether ACSL4 activity induced histone acetylation was associated with the epigenetic regulation of SNAIL expression. In line with the altered histone H3 acetylation, we found a significant decrease in SNAIL expression in *ACSL4* knockdown cells but a considerable increase in *ACSL4* OE cells compared to controls. We also evaluated SNAIL expression following treatment of MDA-MB-231 cells with PRGL493 and observed that it was significantly downregulated (Figure 7B, top, middle, and bottom panels). Furthermore, we carried out ChIP–qPCR assays to define the underlying mechanism of ACSL4-induced SNAIL expression. To delineate the enrichment of H3 acetylation marks in the *SNAI1* promoter, we design the walking primers spanning −1500 bases upstream to the transcription start site (TSS) by utilizing the Eukaryotic Promoter Database based on *SNAI1* chromosomal location (Figure 7C). The ChIP–qPCR analysis reveals the strong enrichment of H3K9 and H3K27 acetylation marks upstream of -1.5 kb, -1.2 kb, -0.8 kb, and -0.5 kb region in MDA-MB-231 cells with no significant enrichment of H3K23 and H3K14 acetylation marks (Figure 7D). We found that the occupancy of histone acetylation (H3K9ac and H3K27ac) marks at the promoter region of *SNAI1* was drastically reduced by *ACSL4* knockdown but was significantly rescued with the exogenous addition of acetyl-CoA (Figure 7E and 7F). Moreover, we observed a markedly enhanced H3K9 and H3K27 acetylation level at *SNAI1* promoter in *ACSL4* OE cells compared to control cells (Figure 7G and 7H). These observations collectively support our notion that ACSL4 activity directs epigenetic regulation of SNAIL expression via histone H3 acetylation.

**Figure 7.**
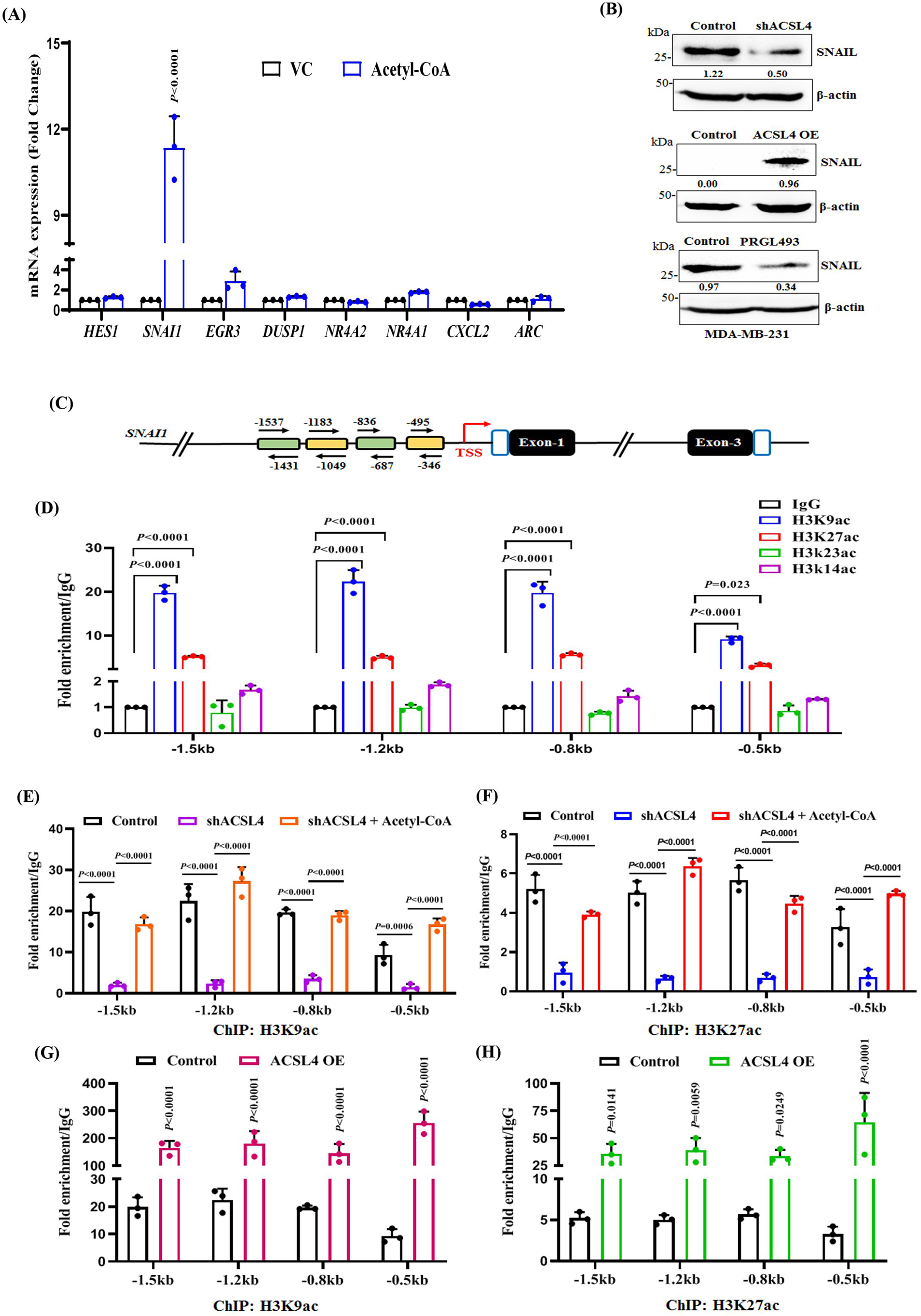
ACSL4 epigenetically regulates *SNAI1* expression via modulating histone H3 acetylation. **A.** The genes that exhibited significant alterations in the RNA-seq analysis were examined by qPCR after supplementation of acetyl-CoA in MDA-MB-231 cells with *ACSL4* knockdown. **B.** Immunoblot analysis of SNAIL expression in *ACSL4* knockdown (top panel), *ACSL4* OE (middle panel), and after treatment with vehicle control or PRGL493 (bottom panel) in MDA-MB231 cells. β-actin was used as a loading control. The densitometric quantifications are shown at the bottom of each immunoblot. **C.** A diagram illustrating the different sites examined for the enrichment of histone acetylation marks at *SNAI1* promoter. **D.** ChIP-qPCR data depicting the enrichment of H3K9ac, H3K27ac, H3K23ac, and H3K14ac using walking primer pairs targeting -1.5 kb to TSS of the *SNAI1* promoter. **E-H**. ChIP-qPCR data showing the enrichment of H3K9ac and H3K27ac at the *SNAI1* promoter, respectively. **E** and **F.** In control, *ACSL4* knockdown, and after supplementation of acetyl-CoA in *ACSL4* knockdown MDA-MB-231 cells. **G** and **H**. In *ACSL4* OE MDA-MB-231 cells compared to control. **D-H,** Each bar represents a mean of triplicate readings of fold enrichment normalized to IgG, error bars, ±S.D. For panels, **A, G, H** Two-way Anova Sidak’s multiple comparisons test, **D** Two-way Anova Dunnett’s multiple comparisons test, and **E, F** Two-way Anova Tukey’s multiple comparisons test were applied.

### ACSL4-SNAIL axis elicits TNBC migration and distant metastasis

ACSL4 mediated regulation of *SNAI1* expression in TNBC encouraged us to explore the role of this regulatory axis in TNBC migration, invasion and metastasis. First, we employed targeted shRNA sequences to meticulously silence the expression of *SNAI1* in ACSL4 overexpressing MDA-MB-231 cells and ascertained the efficacy of this gene knockdown (Figure 8A). Subsequently, in our wound healing assay, we observed that cells with reduced SNAIL expression exhibited impaired wound closure compared to the control cells. (Figure 8B). Likewise, in the trans-well chamber assay, we noted that cells with *SNAI1* knockdown displayed decreased invasion capabilities when compared to the control cells. (Figure 8C). Next, to address the role of ACSL4-H3Kac-SNAIL regulatory axis in TNBC distant metastasis, we isolated MDA-MB-231 cells after orthotopic inoculation in the mammary fat pad of female NSG mice from primary tumor, as well as lung and lymph node metastatic sites. Subsequently, we employed confocal microscopy to analyse the co-expression two proteins at a time in isolated tumor cells. As shown in (Figure 8D and 8E), there was a substantial increase in expression of both ACSL4, H3K9ac, and H3K27ac in lung and lymph node metastatic cells compared to primary tumor cells. We also observed an elevated expression of both ACSL4 and SNAIL in lung and lymph node metastatic cells compared to primary tumor cells (Figure 8F). These observations collectively indicate that SNAIL is essential for ACSL4-mediated TNBC migration/invasion and an appropriate preclinical tumor model at basal state displays a marked correlation between ACSL4-H3K27ac/H3K9ac-SNAIL axes in course of TNBC metastasis.

**Figure 8.**
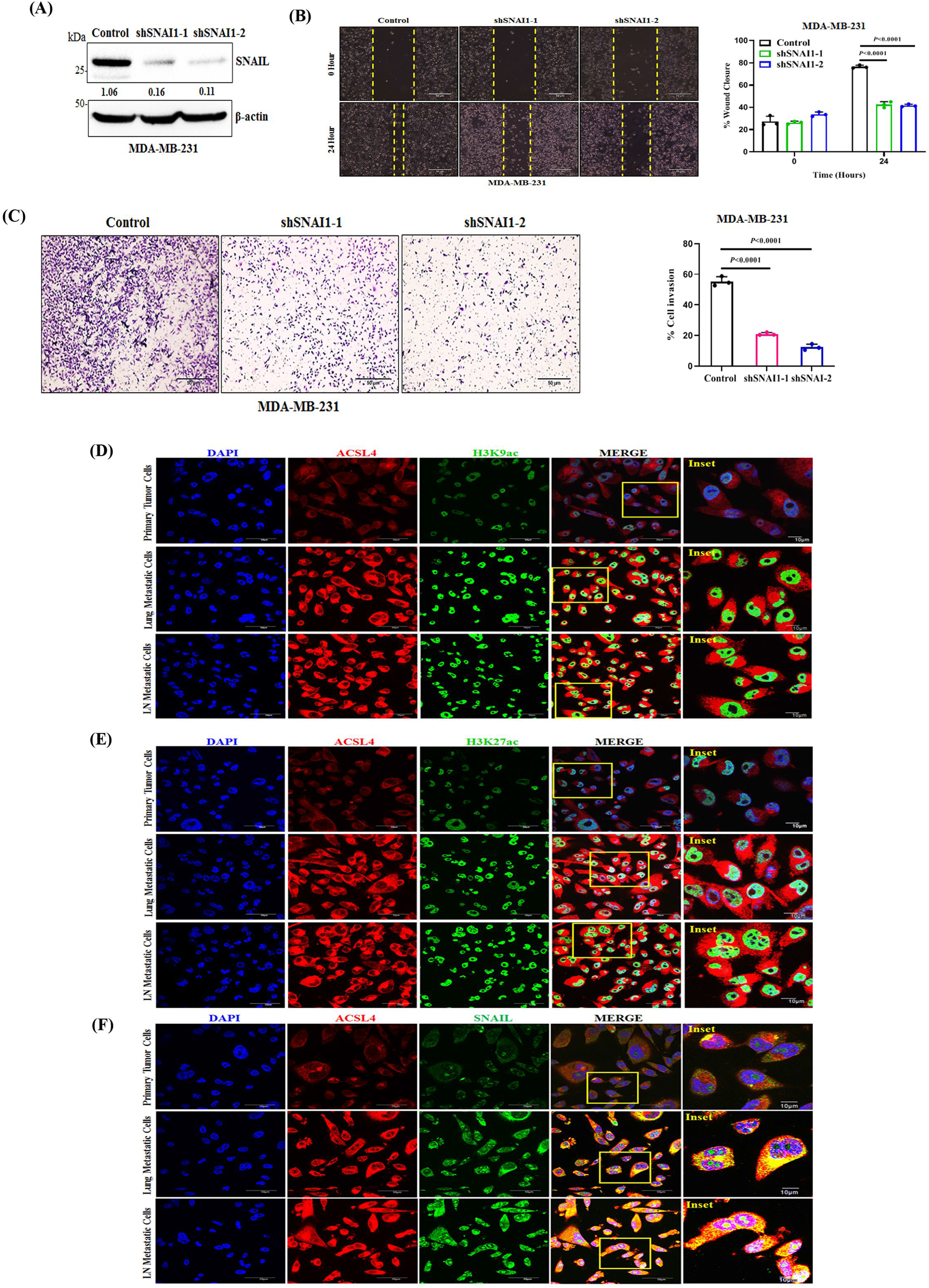
ACSL4-SNAIL axis significantly contributes to the migratory potential of TNBC. **A.** Immunoblot analysis to confirm *SNAI1* knockdown by using two different shRNA in MDA-MB231 cells**. B.** Representative images of the wound healing assay to assess the migration ability of *SNAI1* knockdown MDA-MB-231 cells (left panel), magnification 10×. Scale bar 50 µm. The quantitative bar graphs of the wound healing assay is shown (right panel), error bars, mean ± S.D. **C.** Representative images of the trans-well chamber invasion assay to measure the invasive ability of *SNAI1* knockdown MDA-MB-231 cells (left panel), magnification 10×. Scale bar 50 µm. The quantitative bar graphs of the invasion assay is shown (right panel), error bars, mean ± S.D. **D-F.** MDA-MB-231 cells isolated from primary tumor, metastatic lung, lymph node and subjected to confocal analysis after having either co-staining **(D)** ACSL4 (red) and H3K9ac (green), or (**E)** ACSL4 (red) and H3K27ac (green), or (**F)** ACSL4 (red) and SNAIL (green). In all cases, DAPI was used to stain the nuclei. Scale bar 50 and 10 μm. For panels **B** Two-way Anova Sidak’s multiple comparisons test and **C** One-way Anova Dunnett’s multiple comparisons test were applied.

### ACSL4 and SNAIL expression are markedly upregulated in human TNBC metastasis

To establish a clinical correlation between ACSL4 and SNAIL expression, we mined the TCGA dataset, encompassing breast cancer patients, which is publicly accessible through UCSC Xena Browser. Our findings unveiled a noteworthy upregulation of both *ACSL4* and *SNAI1* mRNA expression levels exclusively within the basal subtype, which is characterized as triple-negative breast cancer (TNBC), when compared with the other subtypes of breast cancer (Figure 9A). In addition, the study of 293 human TNBC samples from public RNA-seq data as analysed by the Breast Cancer Gene-Expression Miner v5.0 (bc-GenExMiner v5.0) showed a positive correlation (*P* = 0.0002, *r* = 0.22) between *ACSL4* and *SNAI* mRNA levels (Figure 9B). Next, to investigate the potential correlation between ACSL4 and SNAIL expression in human TNBC distant metastasis, we conducted immunohistochemical staining on parallel sections of TNBC primary tumors and TNBC distant metastatic (Lung and Lymph node) tissues. The immunohistochemistry analysis of TNBC primary tumors revealed a moderate in situ expression for both ACSL4 and SNAIL proteins (Figure 9C-9E). In contrast, we found marked overexpression of both ACSL4 and SNAIL proteins in TNBC metastatic sites, such as lymph nodes and lungs, as compared to the expression of both of these proteins in TNBC primary tumors. Overall, our in vitro, in vivo, and human patient sample data underscore the potential importance of ACSL4-SNAIL nexus in the metastatic progression of TNBC.

**Figure 9.**
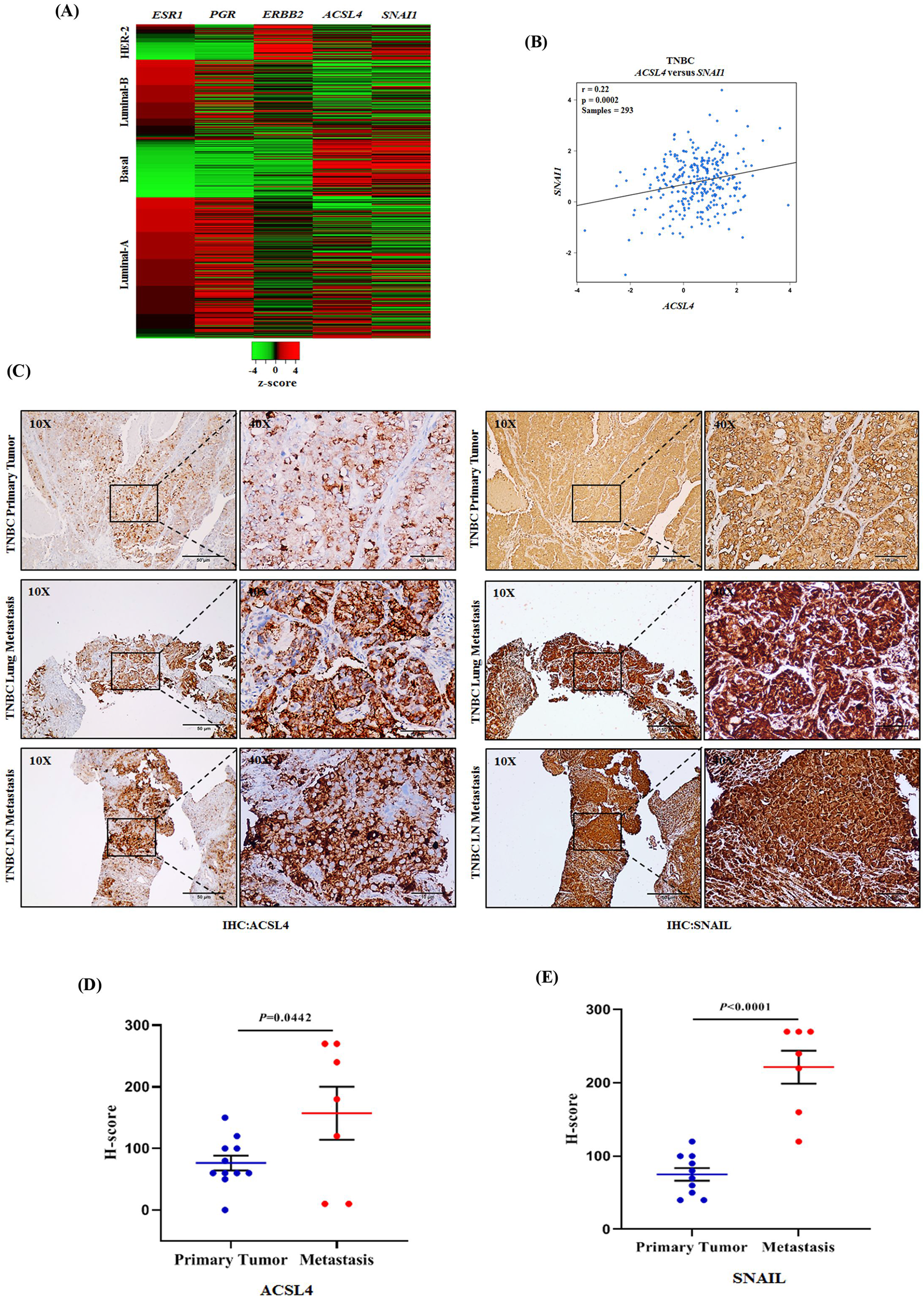
ACSL4 and SNAIL expression are significantly elevated in human TNBC metastasis. **A.** A heat map representation of *ACSL4* and *SNAI1* transcript levels in relation to *ESR1*, *PGR*, and *ERBB2* expression in distinct breast cancer subtypes. Data retrieved from the TCGA Breast Cancer (BRCA) dataset, which is publicly accessible via UCSC Xena Browser. **B.** Scatter plot showing the positive correlation between *ACSL4* and *SNAI1* mRNA expression in TNBC samples as analysed by the bc-GenExMiner v5.0 web tool. Pearson pairwise correlation coefficient on 293 TNBC samples shows a positive correlation (r = 0.22) between *ACSL4* and *SNAI1* and is significantly associated with each other (*p*= 0.0002). **C.** Immunohistochemical staining of formalin-fixed paraffin embedded sections of human TNBC primary tumors and TNBC metastatic tissues using anti-ACSL4 and anti-SNAIL antibodies. Representative photomicrographs were shown at 10X and 40X magnifications (inset). Scale bar, 50 μm (10×) or 10 μm (40×). **D** and **E** Quantitative H-scores for TNBC primary tumors (*n* =11) and TNBC metastatic tissues (*n* =7) were calculated for ACSL4 (**D**)and SNAIL (**E**) expression and represented as dot plots; error bar, mean ±SEM. For panels **D** and **E** Student *t*-test (two-tailed) were applied.

## Discussion

Cancer metabolism has been a burgeoning field over the past two decades, owing to the pivotal link between oncogenes and metabolic pathways ^53^. While the impact of metabolic reprogramming on the metastatic process is poorly understood, accumulating data suggest that FAO is a critical driver of cancer progression and dissemination, with ACSLs emerging as crucial modulators implicated in tumor growth, survival and metastatic spread ^21, 23, 25, 54^. Given, the emerging attention to FAO associated gene signature, which is intricately related to increased aggressiveness, and poor clinical prognosis in breast cancer ^55^, there remains much to be explored regarding the association between deregulated FAO and precise control of TNBC metastasis. Herein, we demonstrate a compelling evidence of a significant correlation between elevated levels of ACSL4 and TNBC, primarily driven by the distinct absence of progesterone receptor (PR). Moreover, the loss of ACSL4 activity profoundly attenuates the metastatic propensity of TNBC’s. Transcriptome analyses and subsequent validation led us to ascertain ACSL4 as a key modulator of TNBC migration and metastasis by altering SNAIL expression. In the realm of mechanistic understanding, ACSL4 influences FAO and acetyl-CoA levels in TNBC, which culminates in the modulation of histone acetylation patterns, and the nuanced epigenetic governance of SNAIL expression. Importantly, we observed a strong correlation between ACSL4 and SNAIL expression in clinical TNBC metastatic samples. Together, our findings provide novel molecular understanding of the complex relationship between metabolic shifts and epigenetic modifications that work together to coordinate metastasis of TNBC.

Recent research by Dattilo et al., has shed light on the regulatory processes governing the differential expression of ACSL4 in breast cancer, specifically highlighting the role of ERRα as a transcription factor responsible for activating the human ACSL4 promoter in TNBC ^56^. While ACSL4’s involvement in promoting an aggressive phenotype in breast cancer is widely acknowledged ^24, 44, 56, 57^, the mechanisms behind its overexpression in TNBC have remained largely unexplored. Notably, several studies have focused on deciphering the downstream elements influenced by ACSL4, but the cellular mechanisms governing ACSL4 expression in both physiological and pathological conditions have remained enigmatic. In this context, our study has made significant strides by identifying progesterone receptor as a key transcriptional regulator of *ACSL4* in breast cancer and provides rationale insight for heightened expression of ACSL4 particularly in TNBC subtype where PR expression is obscure in clinical settings.

Cancer cells utilize metabolic reprogramming as a fundamental strategy to adapt to nutrient scarcity, facilitating their survival, and uncontrolled growth in a transformed state ^9^. Lipid metabolism has garnered considerable attention in recent days due to its significant impact on tumor growth and metastasis ^54^. We believe that TNBC cells depend on ACSL4 for several reasons. ACSL4 occupies a pivotal position in metabolism because it is crucial for both fatty acid oxidation and lipid synthesis by activating long-chain FA’s ^15^. Our findings show that inhibiting ACSL4 leads to a substantial reduction in FAO, a decrease in cellular acetyl CoA levels, and impaired cell migration, underscoring its importance in TNBC progression. Recent reports have highlighted the involvement of fatty acid transport proteins, CD36, FATP1/SLC27A1, and the enzyme CPT1A (carnitine palmitoyl transferase 1A) in the orchestration of metastatic cancer progression ^12–14^. More recently, Camrada et al., have focused on inhibiting fatty acid oxidation (FAO) components to curb TNBC growth ^48^. Although, several metabolic enzymes have been studied as targets for cancer therapy but the vulnerabilities of distinct tumor types to selective metabolic enzyme inhibitors remain largely unexplored ^58^. The use of etomoxir to inhibit FAO has off target effects ^59^. However, ST1326, which inhibits CPT1A, shows promise in a Myc-inducible mouse lymphoma model ^60^. Targeting TNBC’s reliance on FAO is a promising approach, requiring improved clinical stratification and precise metabolic inhibitor application. In this study, we utilized PRGL493, a selective inhibitor of ACSL4 activity, demonstrates robust anti-metastatic potential in TNBC preclinical models though its effect on primary tumor growth was found to be insignificant. However, loss of ACSL4 protein leads to both inhibition of tumor growth and metastasis. Recently, Castillo et al., demonstrated its efficacy in reducing tumor growth in breast cancer ^47^, though they did not show the status of metastasis in experimental animals. Notably, neither the loss of protein or enzymatic activity changes proliferative capacity of TNBCs in vitro but we observe its growth inhibitory effect in in vivo models suggesting the possible impact of tumor microenvironment in modulating the differential response. Together, our investigation unveiled its remarkable ability to effectively suppress FAO and restrain metastasis in TNBC. This makes it critically important for the advancement of ACSL4 inhibitors to the clinics, accompanied by robust medicinal chemistry, structural biology, and adequate pharmacokinetics and pharmacodynamics studies considering the limitations of its current application as possible therapeutics.

The alterations in histone modifications play important roles in transcriptional regulation in cancer cells ^61^. H3K9ac and H3K27ac are crucial epigenetic marks of active transcription are found to be altered in various cancer. H3K9 hyperacetylation is associated with specific gene regulation in breast cancer ^28^. Similarly, H3K27ac acetylation is upregulated in colorectal cancer ^62^. In this study, we elucidate acetyl-CoA as an epigenetic regulating metabolite to enhance TNBC migration and invasion. The regulation of cell migration, epithelial to mesenchymal transition (EMT), and metastasis has been linked to acetyl-CoA production and subsequent histone acetylation. High acetyl-CoA abundance has been associated with elevated H3K27 acetylation at genes associated with cell adhesion and migration in human glioblastoma ^63^. Acetyl-CoA hydrolase (ACOT12) mediated changes in acetyl-CoA levels have been identified to drive hepatocellular carcinoma metastasis by epigenetic induction of EMT ^64^. The depletion of acetyl-CoA contributed by loss in ACSL4 activity and reduced FAO, decreases histone acetylation primarily at H3K9 and H3K27 sites. Furthermore, our study elucidates that ACSL4 activity drives epigenetic regulation through acetyl-CoA-induced histone acetylation in the *SNAI1* promoter region, leading to the activation of SNAIL expression in TNBC. Our intensive investigation of the publicly available transcriptomic datasets of human TNBC patient samples exhibits a significant positive correlation between *ACSL4* and *SNAI1* expression.

Underpinning our observations, recently, Loo et al., revealed that perturbations to fatty acid oxidation by retinoids, promote a more epithelial cell phenotype, blocking EMT-driven breast cancer metastasis through regulation of cellular plasticity ^65^. The link between FAO and tumor progression has been strengthened by studies in which Myc-overexpressing breast cancer exhibited bioenergetic reliance on FAO for growth, and the tumorigenic potential of TNBC cells was dependent on FAO-driven Src activation ^48, 66^. Digging deeper, we amassed several lines of evidence that ACSL4 is the critical ACSL enzyme required not only for the increased migratory potential of TNBC cells in culture but also for the ability of TNBC to metastasize in vivo. Emerging reports suggests that metastatic cancer cells prioritize different metabolic programs distinct from primary tumor in order to enhance their survival throughout the metastatic process, ultimately enabling successful colonization ^67^. Furthermore, FAO has been identified as a critical metabolic adaptation for cancer metastasis to lymph nodes ^68, 69^. Similarly, we also observed a substantial elevated expression of ACSL4, accompanied by increased expression of H3K9ac, H3K27ac, and SNAIL in TNBC cells isolated from metastatic organs such as lung and lymph node than the cells isolated from primary tumor. Our findings also highlight the dynamic changes in ACSL4, H3K9ac, H3K27ac and SNAIL expression during distant metastasis compared to primary tumors. This suggests that ACSL4 mediated modulation of histone acetylation and subsequent regulation of SNAIL expression are crucial events in the metastatic cascade of TNBC. These findings provide valuable insights into the molecular events that drive TNBC metastasis and may open new avenues for therapeutic intervention.

In conclusion, our study uncovers a novel regulatory axis involving ACSL4, H3K9ac, H3K27ac, and SNAIL in TNBC metastasis (Figure 10). The elevated expression of ACSL4 in TNBC, due to selective absence of PR, contributes to enhanced metastatic potential. Targeting ACSL4 or its downstream effectors such as histone acetylation marks and SNAIL could hold promise as therapeutic strategies for combating TNBC metastasis. This research advances our understanding of the molecular mechanisms underlying TNBC aggressiveness and provides potential targets for precision therapeutic approaches in the treatment of this challenging disease.

**Figure 10.**
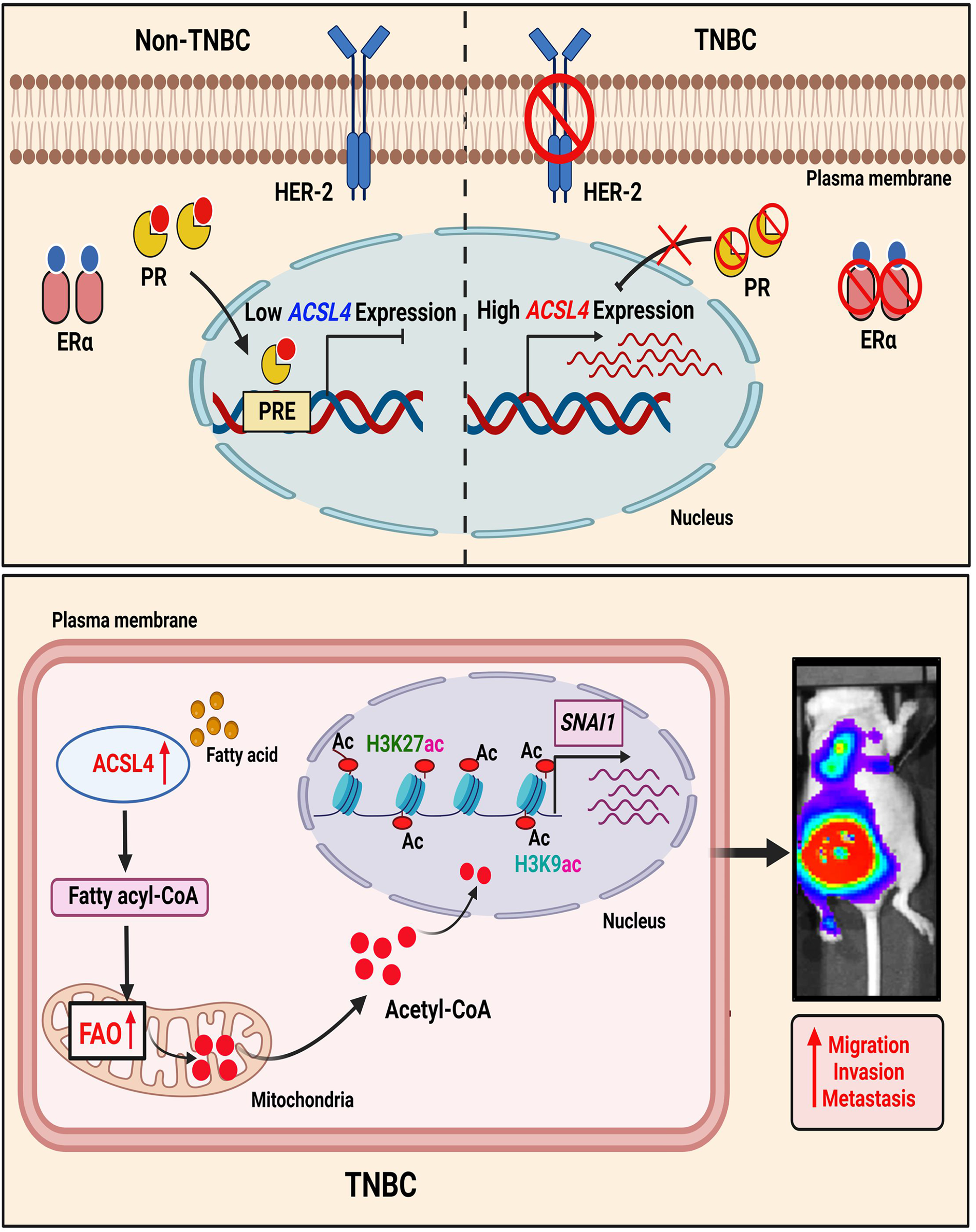
Schematic diagram illustrating the selective absence of progesterone receptor (PR) leads to elevated ACSL4 expression in TNBC, promoting metastasis via ACSL4-driven histone H3 acetylation and SNAIL expression. Illustration depicting the differential expression of *ACSL4* between Non-TNBC and TNBC molecular breast cancer types is a result of progesterone receptor (PR)-driven transcriptional regulation. In the milieu of TNBC, heightened ACSL4 activity plays a pivotal role in fostering elevated FAO and cellular acetyl-CoA levels. This, in turn, leads to enhanced enrichment of H3K9ac and H3K27ac on the *SNAI1* promoter, resulting in elevated SNAIL expression. Consequently, this molecular cascade drives increased invasion, migration, and metastasis in TNBC.

## Methods

### Reagents and antibodies

Dimethyl Sulfoxide (DMSO), bovine serum albumin (BSA), anti-β-Actin (cat# A3854, 1:10,000) antibody, crystal violet dye, and polybrene were purchased from Sigma-Aldrich. RIPA lysis buffer (Pierce^TM^, cat# 89901), protease and phosphatase inhibitor cocktail (100X) (Halt^TM^, cat# 78441) were purchased from Thermo Scientific. PRGL493 was purchased from MedChemExpress (cat# HY-139180). Mifepristone (RU486) was obtained from Sigma-Aldrich (cat# M8046). XenoLight D-Luciferin potassium salt (P/N 122799) obtained from Perkin Elmer. SlowFade™ Gold Antifade Mountant was purchased from Invitrogen (cat# S36938). Isoflurane (FORANE) was obtained from Baxter U.S. Health Care. Matrigel Invasion Chamber (24 well plate 0.8 microns, Lot-8351001) was purchased from Corning. CUT&RUN Assay Kit for ChIP (cat# 86652) and antibodies for ERα (cat# 8644), PR-A/B (cat# 8757), HER-2 (cat# 2165), H3K9ac (cat# 9649), H3K27ac (cat# 8173), H3K14ac (cat# 7627), H3K18ac (cat# 13998), H3K56ac (cat# 4243), and total H3 (cat# 4499) were obtained from Cell Signaling Technology, while H3K23ac was acquired from Abcam (cat# ab177275) and used in 1:1000 dilution for western blot (WB) studies and 1:50 for ChIP studies wherever applicable. ACSL4 antibody (cat# sc-271800, 1:500) was procured from Santa Cruz Biotechnology and Abcam (cat# ab205199, 1:1000). SNAIL antibody (cat# 3879, 1:1000) and (cat# A5243) were purchased from Cell Signaling Technology and ABclonal, respectively. The antibody for GAPDH (cat# 25778, 1:1000) was purchased from Santa Cruz Biotechnology. BCA protein estimation kit, Lipofectamine-3000, FBS (Gibco), RPMI-1640 or DMEM media, Anti-Anti, Puromycin, Alexa Fluor 488/594 conjugated secondary antibodies, and Super Signal West Pico and Femto chemiluminescent substrate were purchased from Thermo Fischer Scientific. PVDF membrane, blocking buffer, iScript Adv cDNA synthesis kit (cat# 1725038), and Clarity Western ECL Substrate (cat# 1705061) were purchased from BIO-RAD. A dual luciferase assay kit (cat# E1910) was purchased from Promega. RNeasy Mini Kit (cat#74104) and QIAprep Spin Miniprep Kit (cat# 27106) were bought from Qiagen. The scrather was purchased from SPL life sciences. The collagenase Type I (cat# 1700-017) was procured from Gibco. Primers for qPCR and ChIP assay were purchased from *Eurofins Genomics, India*. (Details of primer are listed in Supplementary Tables 1-4). All chemicals and antibodies were obtained from Sigma, Cell Signaling Technology, or Thermo Fischer Scientific unless specified otherwise.

### Cell culture

Human breast cancer cell lines: MDA-MB-231, BT-549, HCC-1395, MDA-MB-468, MDA-MB-436, HCC-38, ZR-75-1, MCF-7, and mouse 4T-1-CRL-2539 cell lines were obtained from American Type Culture Collection (ATCC). The 4T-1-Luc-GFP cell was procured from PerkinElmer. Mycoplasma-free early passage cells were revived from liquid nitrogen vapor stocks and inspected microscopically for stable phenotype before use. The cells were cultured in RPMI-1640 or DMEM medium (Gibco™) supplemented with 10% fetal bovine serum (FBS, Gibco™) and anti-anti (Invitrogen, containing 100 μg/mL streptomycin, 100 unit/mL penicillin, and 0.25 μg/mL amphotericin B). The cell lines used in the study are authenticated by STR profiling. The cells were incubated in an Eppendorf Galaxy 170R/170S CO_2_ incubator to provide a stable and homogeneous 5% CO_2_ and 37°C temperature and humid atmosphere required for cell culture.

### Generation of stable cell lines

To generate stable overexpression cells for *ESR1*, *PGR*, and *ACSL4* their respective ORF were amplified from pEGFP-C1-ERalpha (Addgene, cat# 28230), pCDNA3-PRB (Addgene, cat# 89130), and FACL4 (ACSL4) (NM_004458) Human Tagged ORF Clone (cat# RC205356) plasmids and cloned into a 3^rd^ generation lentiviral vector PLJM1-EGFP (Addgene, cat# 19319). The control and *ERBB2* OE cells were generated using retroviral vectors pBABE-puro (Addgene, cat# 1764) and pBABEpuro-ERBB2 (Addgene, cat# 40978), respectively. The 3rd generation lentiviral vector pUltra-Chili-Luc (Addgene, cat# 48688) with the bi-cistronic expression of tdTomato and luciferase was used to create MDA-MB-231 fluorescent-tagged cells. *ESR1*, *PGR*, *ERBB2*, *ACSL4* (mouse), *ACSL4* (human), and *SNAI1* knockdown cells were generated by cloning respective shRNA sequences into the 3rd generation lentiviral plasmid pLKO.1 TRC cloning vector (Addgene, cat# 10878) between unique AgeI and EcoRI sites downstream of the U6 promoter. For the generation of *SNAI1* knockdown, the puromycin resistant gene present in the pLKO.1 TRC cloning vector was replaced with hygromycin resistance. Lentiviral and retroviral particles were generated in HEK293T and Phoenix cells, respectively. Lipofectamine-3000 was used as a transfection reagent. The media containing the viral particles was supplemented with polybrene (8 μg/mL) for transduction. Cells were subjected to puromycin or hygromycin selection as required 48 h post-transduction. The knockdown and overexpression profiles were confirmed after 1 week of selection via Western blot. The list of overexpression primers and shRNA sequences are listed in Supplementary Tables 1 and 2.

### Human samples and ethics approval

Human tissue samples of Non-TNBC and TNBC were provided by King George’s Medical University (KGMU), Lucknow, India. Written informed consent was obtained from above patients with the protocol approved by the Ethics Committee of KGMU (Protocol Number: 116^th^ ECM ІIA/P16) and CSIR-CDRI (Protocol Number: CDRI/IEC/2023/A2). All the research was carried out according to the provisions of the Declaration of Helsinki of 1975. None of these patients had received any chemotherapy or radiotherapy before sample collection. No patients were compensated for participation in this study. Paraffin-embedded tissue sections of human TNBC metastasis were obtained from Rajiv Gandhi Cancer Institute and Research Center (RGCIRC), New Delhi, India. The study protocol was approved by the Institute’s Ethics Board (Protocol Number: Res/BR/TRB-24/2021/43).

### Immunohistochemistry (IHC) staining and analysis

IHC for TNBC patient specimens were performed using Ventana Benchmark XT automated closed system following their recommended protocol. ACSL4 and SNAIL antibodies were purchased from Abcam and ABclonal, respectively. All stained slides were examined under EVOS XL core microscope under 10X and 40X magnification. The staining intensity of each section was scored as described previously ^29^ where 0 (no staining), 1+ (weak staining), 2+ (moderate staining), or 3+ (strong staining). The tumor cell positive rate (0–100%) per slice was multiplied by the staining intensity to get an overall H-scores ranging from 0 to 300.

### Luciferase assay

ACSL4 promoter sequence 1000 bp (-1.0 kb upstream of TSS) were amplified from MDA-MB-231 cell genomic DNA by PCR. The amplified fragments were cloned into the PGL4.12 [luc2 CP] vector between the NheI and XhoІ restriction sites. The control and *PGR* knockdown MCF-7 cells were seeded at 70–80% confluence in 6-well plates and transfected with 2 µg of PGL4-1.0 kb ACSL4-promoter, and 50 ng of PGL4 (hRluc-CMV) plasmid using lipofectamine-3000 as transfection reagent (Invitrogen). For RU486 treatment, the PGL4-PGL4-1.0 kb ACSL4-promoter transfected MCF-7 cells were treated either with vehicle control or RU486 (20 µM) for 36 h. Lysis buffer provided in the Dual-Glo Luciferase assay kit (Promega) was used for cell lysis. The GloMax® 96 Microplate Luminometer was used to measure the activities of Firefly and Renilla luciferases according to the manufacturer’s protocol (Promega). Firefly luciferase activity was normalized to Renilla luciferase activity for each sample. Luciferase ACSL4 promoter primers are enlisted in Supplementary Table 3.

### Western Blotting

The cells were harvested and subjected to lysis using RIPA lysis buffer with phosphatase and protease inhibitor cocktail and incubated at -20°C for 24 h, subsequently the samples were subjected to probe sonication with 3 cycles of 10 seconds on, 10 seconds off. The samples were incubated on ice for 5 min followed by centrifugation at 12000 rpm for 15 min. The supernatant was transferred to a new microcentrifuge tube and stored at -80°C. The protein concentration of the lysates were estimated using BCA protein assay kit (Pierce™). An equal quantity of protein for the respective samples were resolved by SDS-PAGE and transferred to a PVDF membrane (Bio-Rad). The membrane was blocked with 5% nonfat dry milk, followed by incubation with appropriate dilutions (manufacturer’s protocol) of primary antibodies overnight at 4°C. Subsequently, the membrane was incubated with a 1:5000 dilution of horseradish peroxidase-conjugated secondary antibody for 1 h at room temperature. Immunoreactivity was detected by enhanced chemiluminescence solution (Bio-Rad Clarity Western ECL Substrate) and scanned by the gel documentation system (Bio-Rad chemidoc XRS plus). Densitometric analysis was performed with ImageJ™, an open-source software (NIH)^70^.

### RNA-sequencing (RNA-seq) analysis

Illumina NovaSeq 6000 was used to perform RNA-sequencing, followed by the Fast QC and Multi QC tools were used to assess the data quality. Human genome (GRch38) was downloaded from iGenomes and indexed using Bowtie2-build with default parameters. Adapter sequences (P7 adapter read1: AGATCGGAAGAGCACACGTCTGAACTCCAG TCA and P5 adapter read2: AGATCGGAAGAGCGTCGTGTAGGGAAAGAGTGT) were removed using Trim Galore (v 0.4.4) and each of the raw Fastq files were passed through a quality check using FastQC. PCR duplicates were removed using the Samtools 1.3.1 with the help of ‘rmdup’ option. Each of the raw files was then aligned to Human genome assembly using TopHat2 with default parameters for paired-end sequencing ^71^. Subsequent to aligning, quantification of transcripts was performed using Cufflinks, and then Cuffmerge was used to create merged transcriptome annotation. The differentially expressed (DE) genes were identified using Cuffdiff. The threshold for DE genes was log2 (fold change) > 1.5 for up-regulated genes and log2 (fold change) < 1.5 for down-regulated genes with p-value < 0.05. Finally, Gene Ontology (GO) enrichment analysis was performed using PANTHER overrepresentation test ^72–74^. The Fisher’s exact test was used and only GO terms with a FDR of < 0.05 were considered. The most significant GO terms related to the phenotype observed in our study were represented in a heat map.

### Quantitative real-time PCR

RNA was isolated from cultured cells with RNeasy Mini Kit (Qiagen, cat# 74104), following the extraction protocol suggested by the manufacturer. The concentration and purity of the RNA samples were determined using nanodrop. Reverse transcription (RT) was performed from 2 to 5 µg of RNA with iScript Adv cDNA synthesis kit (Bio-Rad #1725038). The cDNA was diluted 5 times and 1 μL of this was used for each reaction in real-time PCR. Quantitative real-time PCR was carried out using an ABI Step One Plus Real-Time PCR System (Applied Biosystems). Reactions for each sample were performed in triplicate. 18S rRNA amplification was used as the reference gene. The relative expression ratio of target/reference gene was calculated with the ΔΔCt method. For amplification of *ACSL4*, *ACSL1*, *ACSL3*, *ACSL5*, *ACSL6*, *HES1*, *EGR3*, *NR4A1*, *DUSP1*, *ARC*, *SNAI1*, *CXCL2*, and *NR4A2* we performed SYBR Green-based RT-PCR following manufacturer’s instructions. Primers sequences are listed in Supplementary Table 4.

### Chromatin immunoprecipitation (ChIP) assay

ChIP assay was conducted using the CUT&RUN Assay Kit (Cell Signaling Technology, cat# 86652), following the manufacturer’s protocol. Briefly, cells were harvested, washed, and bound to activated Concanavalin A-coated magnetic beads and permeabilized. The bead– cell complex was incubated overnight with the respective antibodies at 4°C. Cells were washed three times and resuspended in 100 μL pAG/MNase and incubated for 1 h at 4°C. The DNA fragments were released into the solution by incubating at 37°C for 10 min. The DNA fragments were purified using the DNA purification buffers and spin columns (ChIP, CUT&RUN, Cell Signaling Technology, cat# 14209) and SYBR Green-based real-time PCR was conducted. Primer sequences used for the ChIP experiments for different genes are enlisted in Supplementary Table 5.

### ACSL activity assay

The ACSL activity was assessed using Acyl-CoA Synthetase (ACS) Assay Kit (Abcam, cat# ab273315). The cells were homogenized in ACS Assay Buffer followed by centrifugation at 10,000x g at 4°C for 15 min. The supernatant was collected and stored at -80°C. Protein was measured using BCA Protein Assay kit (Pierce). Equal concentration of cell lysates were used for the individual group of samples to be analyzed. The volume was adjusted to 50 µL/well with ACS Assay Buffer followed by the addition of 50 µL of reaction mix (ACS Substrate, ACS Enzyme mix, ACS Converter, ACS Developer, and ACS Probe). In the assay, acyl-CoA generated by ACS activity is metabolized by the Enzyme mix, Developer mix, and Converter mix to produce an intermediate compound that reacts with a probe to produce a fluorescence signal that was be detected at Ex/Em = 535 /587 nm.

### Fatty Acid Oxidation (FAO) analysis

To measure cellular FAO using oxygen consumption rate (OCR) assay, 40,000 cells were plated in cell culture microplates and cultured in complete growth medium (RPMI-1640, 10% FBS) overnight at 37°C in 5% CO_2_ incubator to allow cell attachment. The growth medium was exchanged to substrate-limited medium (Seahorse XF DMEM medium (Agilent, cat# 103575-100), 1 mM Seahorse XF pyruvate solution (Agilent, cat# 103578-100), 10 mM Seahorse XF glucose solution (Agilent, cat# 103577-100), 2 mM Seahorse XF glutamine solution (Agilent, cat# 103579-100)) on the day of experiment. The oxygen consumption rate (OCR) assay was performed using the XF long-chain fatty acid oxidation (LCFA) stress test kit (Agilent, cat# 103672-100) and Seahorse XFe24 Analyzer was used to investigate the long-chain fatty acid OCR according to the manufacturer’s protocol.

### Acetyl-CoA estimation assay

The acetyl-CoA content was determined in Acetyl-CoA Assay Kit (Abcam, cat# ab87546). Briefly, acetyl-CoA standard curve was made in the range of 0–100 pM and the correlation coefficient was 0.990 or higher. The protein in the sample was removed using the perchloric acid method and the supernatant was neutralized with 2 M KOH. The CoA Quencher and Quencher remover were added into the sample to correct the background generated by free CoA and succ-CoA. The sample was then diluted with the reaction mix, and the fluorescence signal was measured at Ex/Em = 535/589 nm using a microplate reader.

### Confocal microscopy

The cells were fixed with 4% paraformaldehyde in PBS for 10 min at RT, followed by permeabilization for 10 min with PBS containing 0.1% TritonX-100 (Sigma-Aldrich) and blocking with 5% BSA in PBS for 1 h at RT. After overnight incubation with primary antibodies (anti-ACSL4, anti-H3K9ac, anti-H3K27ac, and anti-SNAIL), the cells were washed twice with PBS and incubated with fluorophore-conjugated secondary antibodies and DAPI for 1 h at RT. The cells were then washed and mounted with the antifade mounting medium on glass slides. The fluorescence confocal images were acquired by using an Olympus BX61-FV1200-MPE microscope.

### Wound healing assay

In the wound healing assay, 500,000 cells were seeded into individual wells of a 6-well plate and allowed to grow overnight until they formed a confluent monolayer. A scratch was then created using a scrather to generate a straight-line gap in this cell layer. Subsequently, the cells were rinsed with PBS and cultured with fresh complete RPMI-1640 media at 37°C for different time intervals. The cells were incubated at 37 °C for different time intervals with or without specific treatments. The percentage of wound-healed area was determined by analyzing the images using ImageJ Software.

### Cell invasion assay

Invasion assays for 4T-1 and MDA-MB-231 cells were performed using BioCoat Matrigel® invasion chambers (Corning). Briefly, cells were seeded in 500 µL serum-free medium onto the upper compartment of the trans-well while the lower compartment was filled with RPMI-1640 medium, supplemented with 10% FBS acting as chemoattractant. After 24 h of incubation at 37 °C, the non-invasive cells were removed with a cotton swab. The cells that invaded through the membrane and adhered to the lower surface of the membrane were fixed with 100% methanol for 10 min. The invasion chambers were stained with crystal violet for 1 h and then washed twice with PBS. Cell invasion was quantified by counting the cell number using ImageJ Software.

### Colony formation assay

The 4T-1 cells or MDA-MB-231 cells were seeded at 100 or 1000 cells per well in 6-well plates respectively. For experiments involving specific treatments, post 24 h, the adhered cells were treated either with the vehicle or PRGL493 (20 µM) and incubated for a week at 37°C. Subsequently, the media was aspirated, followed by two times PBS wash. The washed cells were fixed with ice-cold methanol for 15 min at 4°C and stained with 0.5% crystal violet dye for 1 h. The excess stain was washed with water, and the plates were allowed to air dry. The stained colonies were counted using ImageJ Software.

### Animal studies

All animal studies were conducted by following standard principles and procedures approved by the Institutional Animal Ethics Committee (IAEC) of CSIR-Central Drug Research Institute (Protocol Number: IAEC/2022/22 and IAEC/2023/129). The experimental mice were maintained in IVC cages under pathogen-free conditions with a 12 h light and 12 h dark cycle, a temperature maintained at 24 ± 2°C, and humidity level of 45 ± 5%. These mice were fed with irradiated standard mouse diet and were kept at the CSIR-CDRI Central Laboratory Animal facility. 4-6 week-old female Balb/c nude mice or NOD *scid* gamma (NSG) mice were used for the different experimental studies. For orthotopic inoculation, different tagged (GFP-Luc, Td-Tomato) 4T-1 cells (1 × 10^6^) in 100 μL PBS were injected into the mammary fat pad of 4–6-week-old nude Crl: CD1-Foxn1^nu^ female mice. For xenograft experiments, 5 × 10^6^ MDA-MB-231 Chili-Luc cells were resuspended in 1:1 PBS: matrigel solution and orthotopically inoculated into the mammary fat pad of 4-week-old NOD.Cg-Prkdcscid Il2rgtm1Wjl/SzJ (NSG) female mice. PRGL493 (500 µg/kg dose) or vehicle (10% DMSO + 40% PEG + 5% Tween-80 + 45% PBS) was administered consecutive days by oral gavage up to 25 days for 4T-1 allograft and 56 days for MDA-MB-231 xenograft experiments from 10 days post tumor cell inoculation. Throughout the study, tumor volume (mm^3^) were measured with a digital vernier calliper at regular intervals, and calculated using the standard formula V = (W^2^ × L)/2 (‘W’ is the short and ‘L’ is the long tumor axis). For live animal bioluminescent imaging studies using IVIS spectrum, Perkin Elmer, 1.5 mg/kg D-Luciferin in PBS was intraperitoneally administered in each tumor-bearing mice. Subsequently, mice were anesthetized by isoflurane and images with ventral positions were obtained using a Perkin Elmer IVIS system coupled with bioluminescence image acquisition and analysis software. Using Living Image software, regions of interest (ROI) on the tumor and metastatic sites were detected and quantified as counts per second.

### Isolation and culture of metastatic cells

4T-1 tumor-bearing mice were sacrificed, and different organs (lymph node, lung, and liver) were harvested under sterile conditions. The single-cell suspension was prepared according to the standard protocol. Briefly, chopped tissues were incubated in HBBS solution with 1 mg/ml of collagenase on a shaker for 3-4 h at 37 °C. After centrifugation at 500×g for 5 min and washing, the cell pellet was re-suspended in regular growth medium (RPMI, 10% FBS, 1× anti-anti) in a T-25 flask. Following a 24 h incubation period, fresh media containing 60 μM 6-Thioguanine (6-TG) was added and the cells were cultured for 3 days to select for 6-TG resistant 4T-1 cells. For the isolation of metastatic cells from MDA-MB-231 xenograft mice model, MDA-MB-231 tumor-bearing mice were euthanized and subsequently primary tumors and different organs (lung and lymph node) were aseptically harvested. The puromycin resistant MDA-MB-231 cells were isolated using the same method as described above, and 2 µg/ml puromycin was used for the selection of tumor cells. Isolated cells from primary tumors and metastatic organs were used for further analysis.

### Isolation of tumor cells from peritoneal fluid

For the isolation of tumor cells from peritoneal fluid, the 4T-1 tumor bearing mice were euthanized and the outer skin of the peritoneum was cut open and the skin was gently pulled back to expose the inner skin lining the peritoneal cavity. Subsequently, ice-cold PBS was injected into the peritoneal cavity using a needle and the resultant peritoneal fluid containing a cell suspension was collected and placed the tubes on ice. The cell suspension was then centrifuged at 1500 rpm for 10 min. The supernatant was discarded and the cell pellet was resuspended in the preferred media containing 6-TG.

### Isolation of Circulating Tumor Cells

For the isolation and propagation of circulating tumor cells (CTC’s), the mice were humanely euthanized at experimental end-point, followed by collection of the mice blood (1 mL) using an EDTA tube by heart puncture. The RBC was lysed using 1X RBC lysis buffer (BioLegend cat# 420301) for 10 min at RT. The contents were centrifuged for 5 min at 230×g. After centrifugation, a whitish pellet was obtained and the reddish supernatant was carefully discarded. The whitish pellet was washed twice with PBS and finally suspended in the desired growth media containing 6-TG to select the 4T-1 cells.

### Flow Cytometry

Flow cytometry was used to determine the percentage of Td-tomato-tagged control and untagged *ACSL4* knockdown 4T-1 cells isolated from distinct metastatic sites. Briefly, cells were harvested with TrypLE (Invitrogen) for single-cell suspension in FACS buffer (PBS with 0.1% BSA and 1 mM EDTA). Subsequent to centrifugation and washing, the cell pellets were resuspended in FACS buffer and analyzed by FACS Calibur (BD Biosciences). The acquired data were analyzed using FlowJo software.

### Analysis of the breast cancer dataset

Clinically relevant associations based on mRNA expression levels of major metabolic enzymes and transporters of the long chain fatty acid oxidation (FAO) pathway were studied using the TCGA Breast cancer (BRCA) dataset, available from the UCSC Xena Browser ^75^. Illumina HiSeq mRNA data of patients with breast cancer (TCGA (BRCA) was downloaded from UCSC Xena for *ESR1*, *PGR*, *ERBB2, ACSL4*, and *SNAI1* expression. The breast cancer dataset was segregated into a PAM50 subtype for Luminal A, Luminal B, HER2+, and Basal subtypes, and a heat map was generated. To study the correlation between *ACSL4* and *SNAI1* expression in TNBC patient samples we employed the “correlation”^76^ module of the Breast Cancer Gene-Expression Miner (bc-GenExMiner v5.0) web-tool and represented the data in a scatter plot.

### Statistics and reproducibility

For statistical analyses, one-way ANOVA, two-way ANOVA, Student’s *t*-test, and Unpaired two-tailed Welch’s *t*-test were used for independent experiments or otherwise mentioned in the respective figure legends. Data are presented as mean ± SEM or mean ± SD, as indicated, of at least three independent experiments or biological replicates. These analyses were done using GraphPad Prism software. The results were considered statistically significant for *p*-values ≤ 0.05 between groups.

## Acknowledgments

We sincerely acknowledge the excellent technical help of Mr. A. L. Vishwakarma for Flow Cytometry, Mr. Anil Verma for Confocal Imaging, Mr. Ajay Singh Verma for Seahorse and luciferase assays. We express our deepest gratitude to Dr. SK Rath for providing the Imaging facility. Authors are immensely grateful to Dr. Juhi Tayal, Dr. Anurag Mehta, Dr. DC Doval and Ms. Somika Tiwari (Biorepository, Rajiv Gandhi Cancer Institute and Research Centre, New Delhi, India) for providing TNBC tumor samples. We extend our heartfelt gratitude to Dr. Jayanta Sarkar for his support in procuring immunocompromised animals. Research of all the authors’ laboratories was supported by CSIR Pan Cancer Research Grant (HCP-40 to DD, MPS) and Fellowship grants from CSIR (AS, KKS, SRS), DST (MAK), and UGC (BM, PR). Further, DD acknowledges grant support from CSIR-FTT (MLP-2025), DST (EMR/2016/006935), DBT (BT/AIR0568/PACE-15/18) and ICMR (2019-1350). Diagrammatic Figures were created using BioRender software (https://biorender.com) and DD owns a full license to publish. Institutional (CSIR-CDRI) communication number for this article is 135.

## Author Contributions

AS involved in study designing, performed experiments and wrote the draft manuscript. KKS, MAK, SRS, AV, and MAN helped in carrying out in vitro and in vivo studies. MPS, NB, and MV performed bioinformatic analysis. KT, BM, PR, SM, ATS, KS, AKM carried out IHC studies. DD conceived the idea, designed experiments, analysed data, wrote the manuscript and provided overall supervision. All authors read and approved the final manuscript.

## Supplementary Information

**Supplementary Figure 1.**
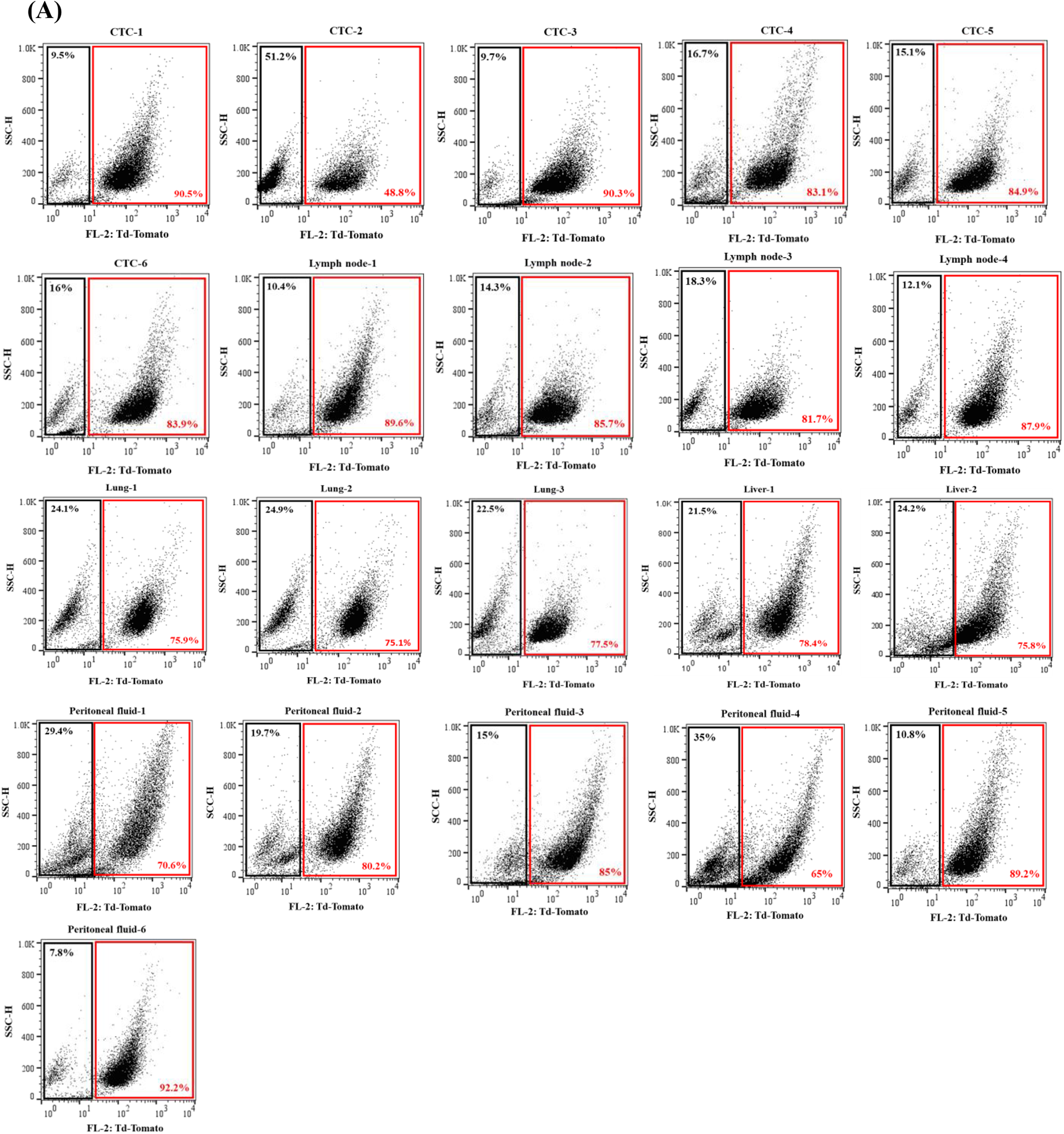
Genetic ablation of *ACSL4* abrogates TNBC distant metastasis. **A.** Representative flow cytometry-derived scatter plots showing control (Td-Tomato-tagged) and *ACSL4* knockdown (untagged) 4T-1 cells isolated from distant metastatic sites (CTC (n=6), lymph node (n=4), lung (n=3), liver (n=2), and peritoneal fluid (n=6)), following orthotpic inoculation in left and right mammary fat pad of 4 to 6-week-old female nude Crl: CD1-Foxn1nude, respectively.

**Supplementary Figure 2.**
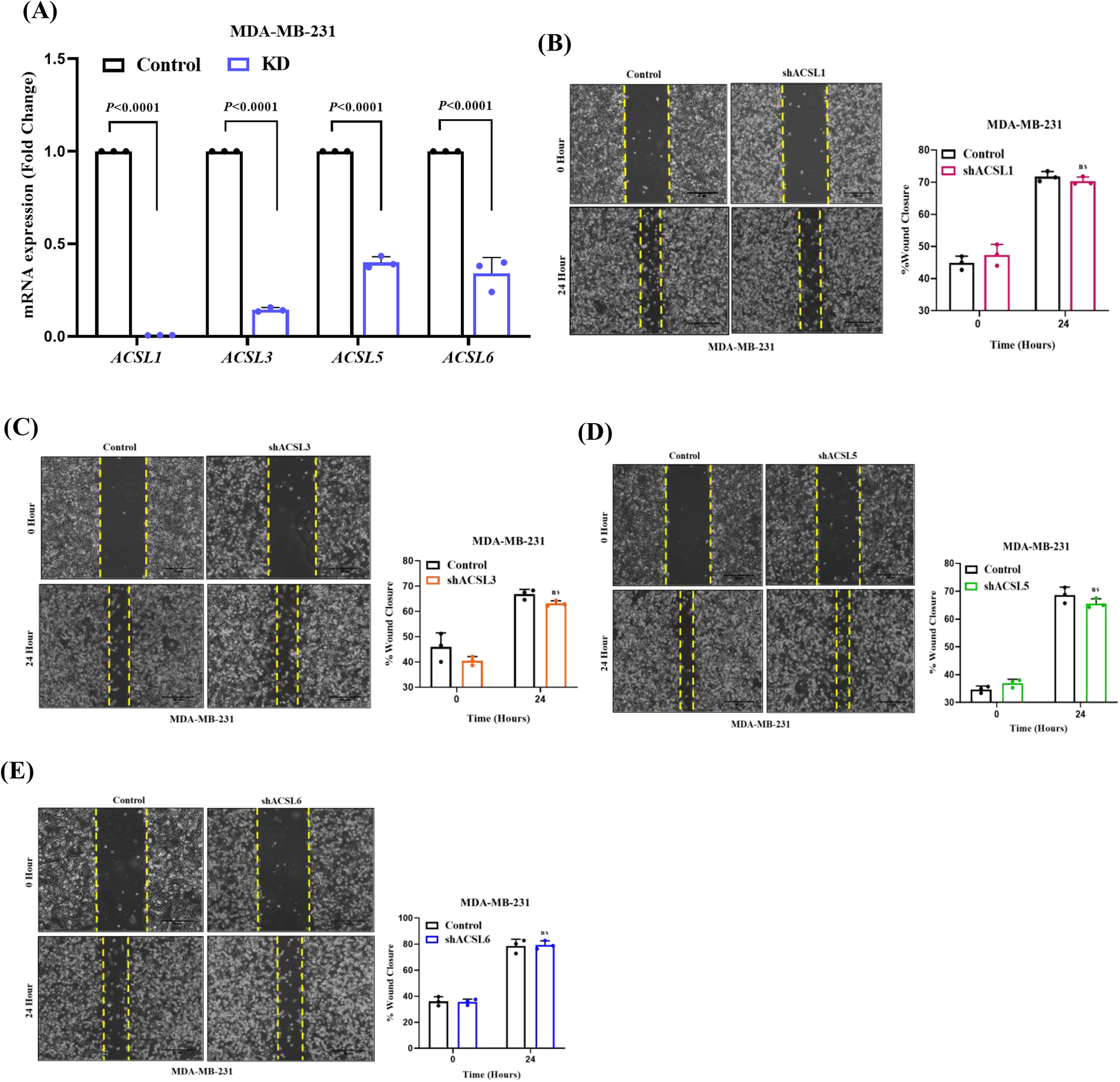
Impact of genetic ablation of *ACSL* isoforms on TNBC migration. **A.** The knockdown of *ACSL* isoforms (*ACSL1*, *ACSL3*, *ACSL5*, and *ACSL6*) using shRNA mediated lentiviral transduction of MDA-MB-231 cells were confirmed by qPCR. The quantitative bar graphs represent mean ± SD. **B-E.** Representative images of wound healing assay in control and *ACSL1* knockdown **(B,** left panel), *ACSL3* knockdown **(C,** left panel), *ACSL5 kn*ockdown **(D,** left panel), and *ACSL6* knockdown (**E**, left panel) in MDA-MB231 cells are shown. 10× magnification. Scale bar 50 µm. The quantitative analysis of wound healing assay (**B, C, D,** and **E,** right panel) are shown. Each bar represents a mean of triplicate readings of samples, error bar, ±SD. For panels, **A, B, C, D,** and **E** Two-way Anova Sidak’s multiple comparisons test was applied.

**Supplementary Table 1.**
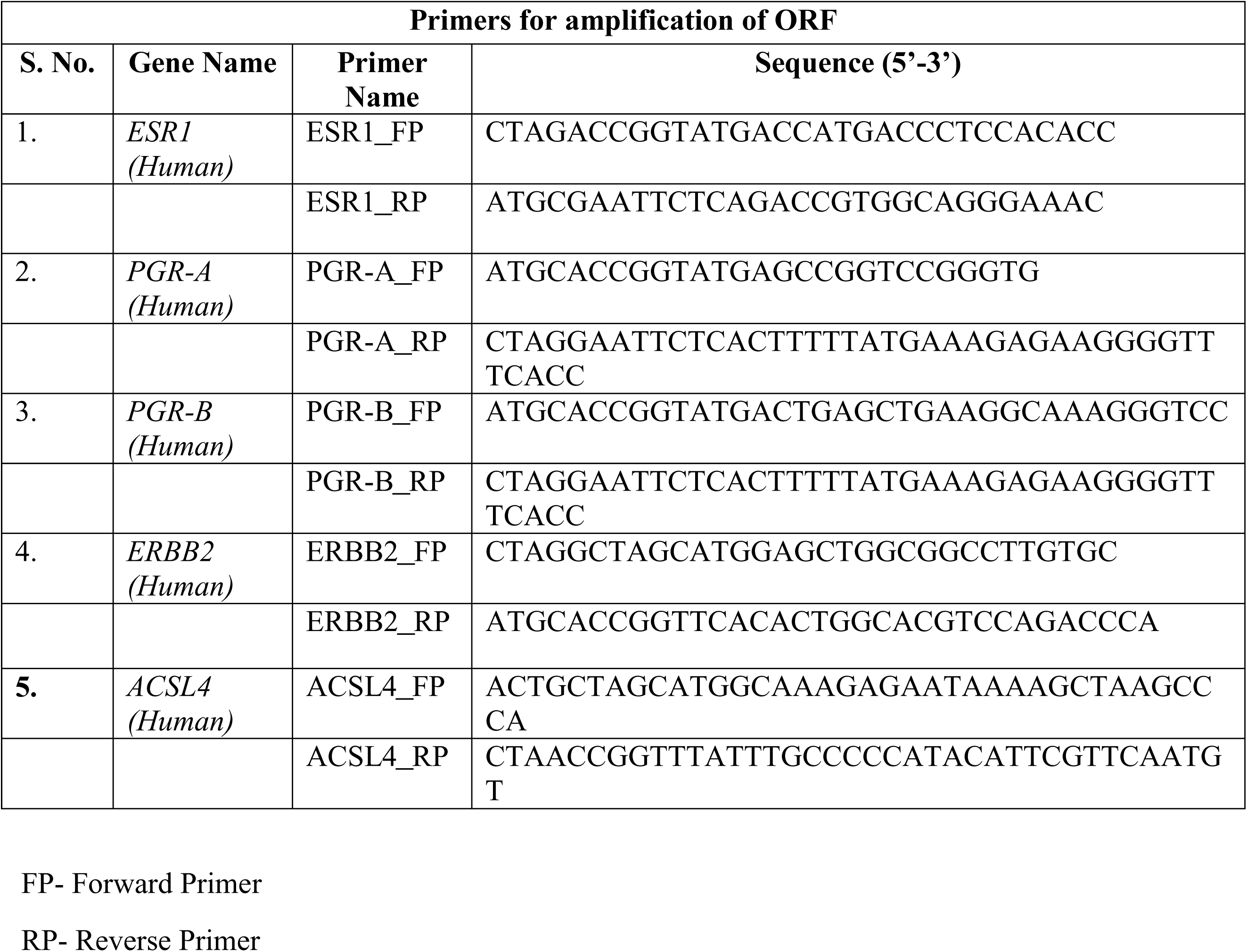
List of overexpression primers.

**Supplementary Table 2.**
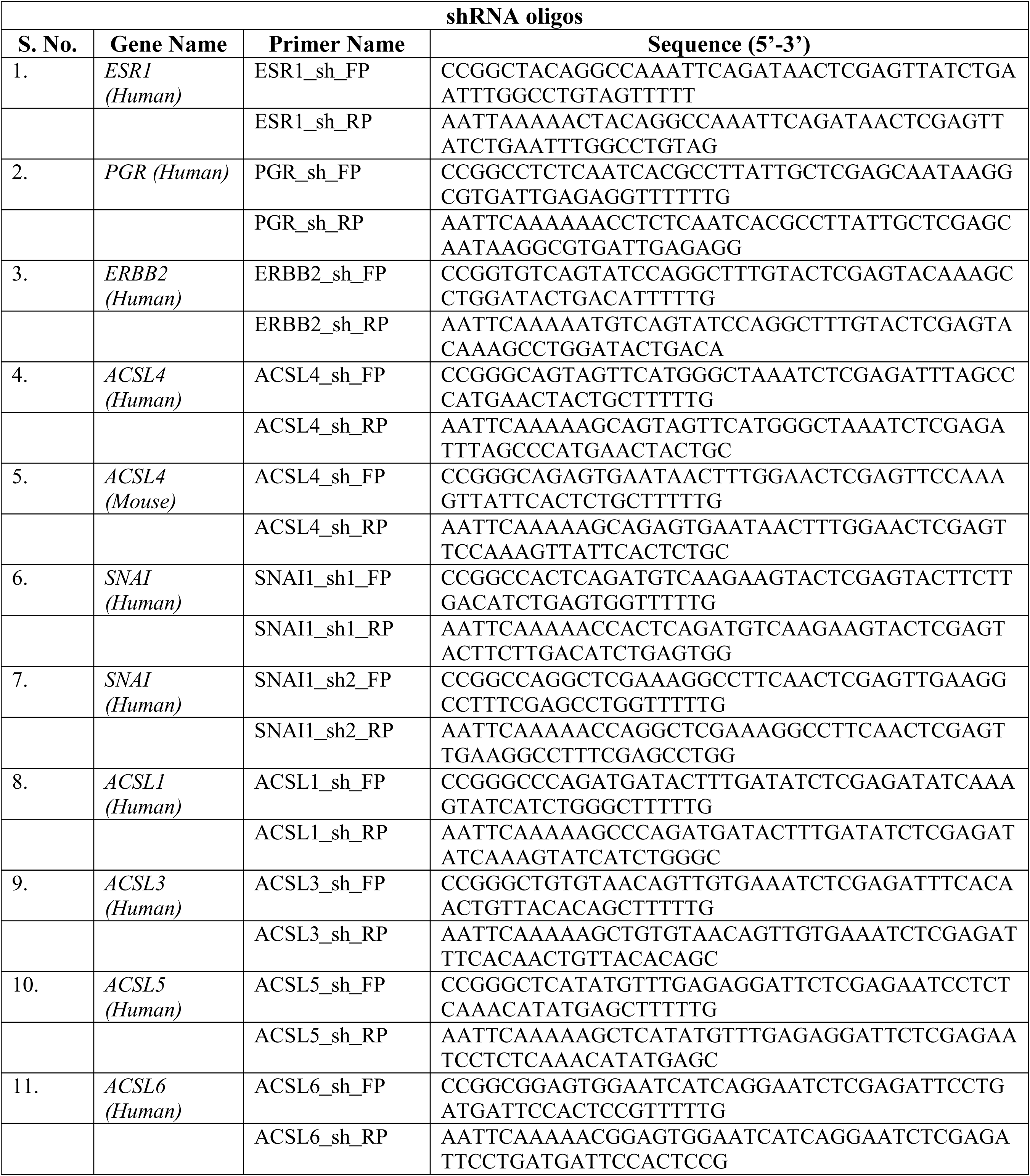
List of shRNA oligos.

**Supplementary Table 3.**
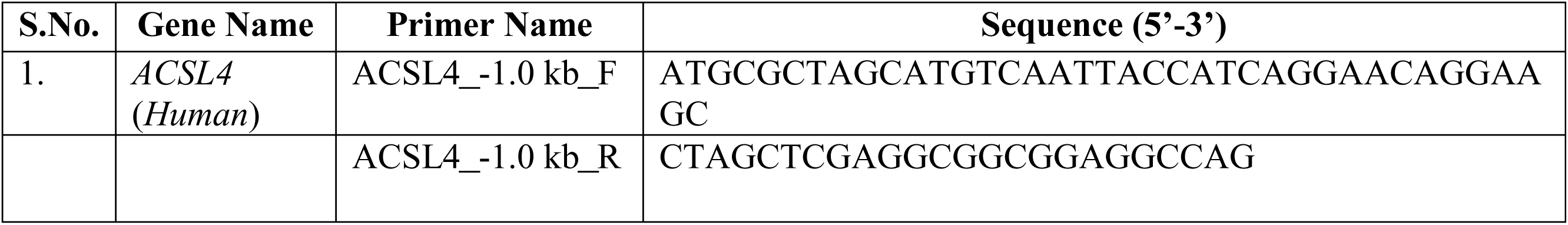
Luciferase ACSL4 promoter primers.

**Supplementary Table 4.**
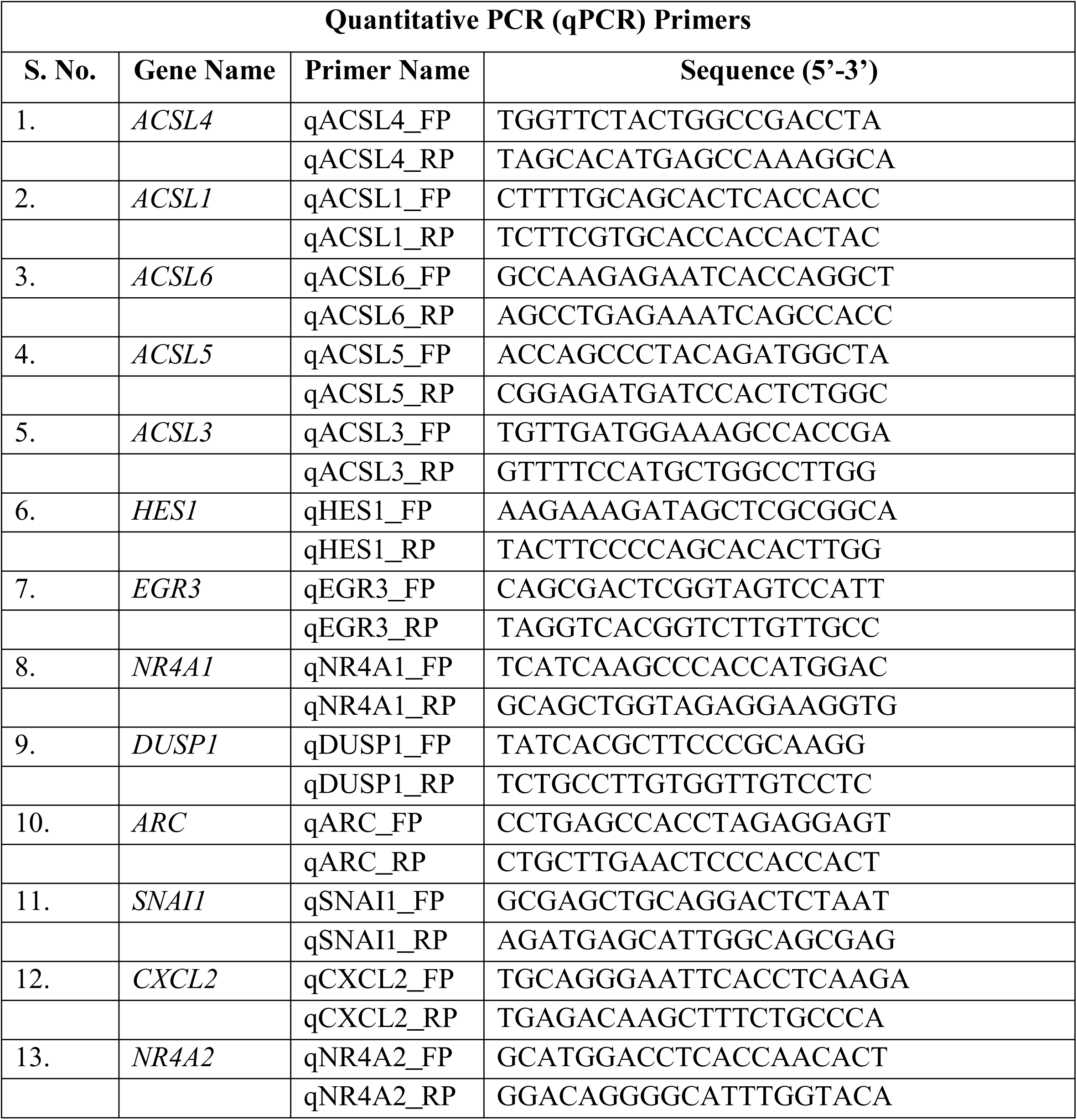
List of qPCR primers.

**Supplementary Table 5.**
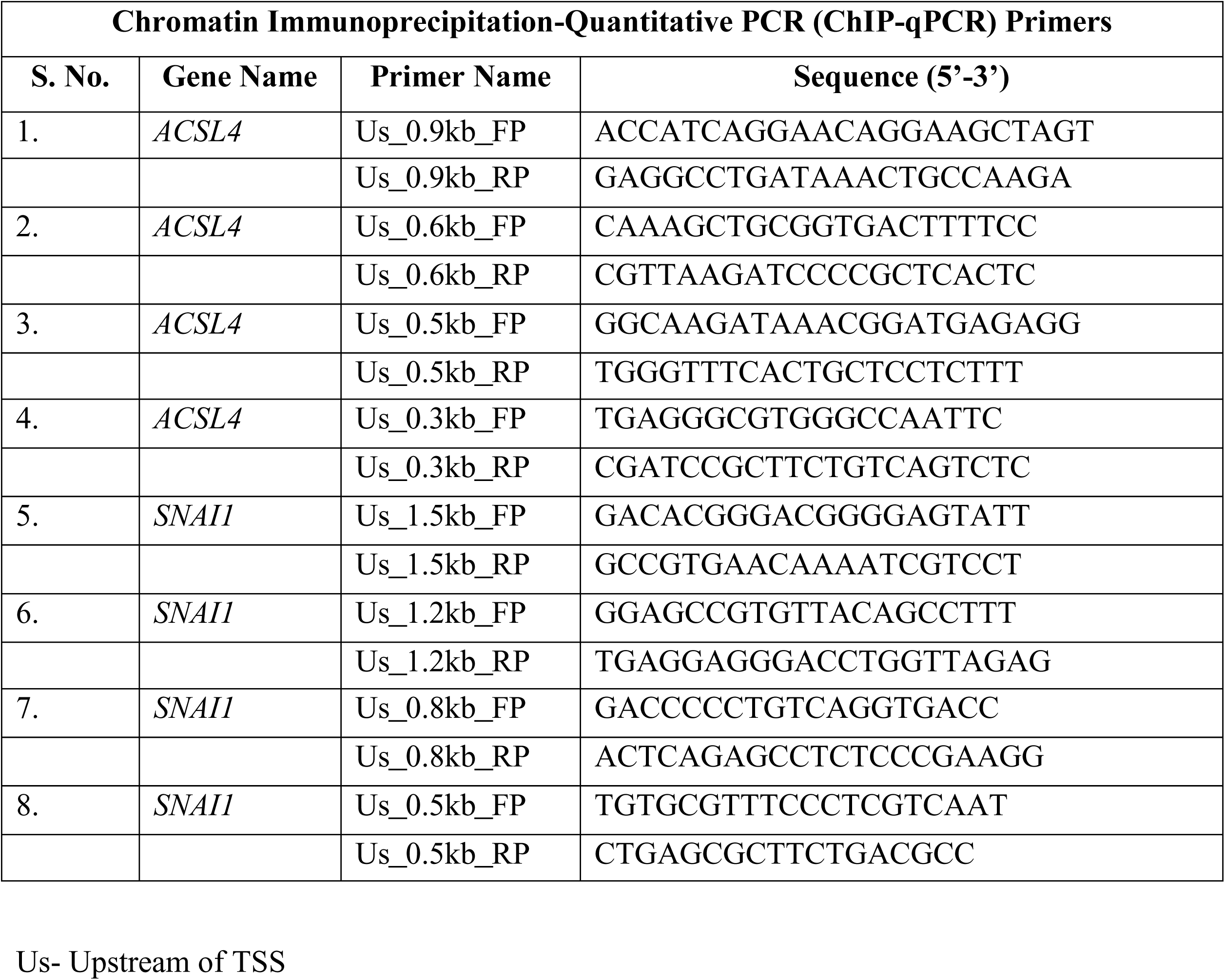
List of ChIP-qPCR primers.

